# Estimating gross transcription rates from RNA level fluctuation data and the effects of sampling time intervals

**DOI:** 10.1101/2023.05.24.541915

**Authors:** Zhongneng Xu, Shuichi Asakawa

## Abstract

Transcription rates are key biological parameters, but the estimation of transcription rates from RNA level fluctuation data by current methods is still problematic, considering in particular the derived relationship between RNA fragments from different samples and the neglect of the effects of sampling time intervals. Based on defining the gross transcription rate as the amount of converted complete nascent RNA divided by time, the present study developed an algorithm that calculated the cumulative transcription amount and RNA abundance at each time point by simulating moving windows to estimate gross transcription rates from RNA level fluctuation data and explore the effects of sampling time intervals on the estimation. The results showed that the gross transcription rates could be calculated from RNA level fluctuation data with the models fitting the experimental data well. In the analysis of 384 yeast genes, the genes with the highest gross transcription rates mainly played roles in cell division regulation and DNA replication, and the gene utilizing the most cellular resources for gene expression during the experiment was YNR016c, whose main functions are fatty acid biosynthesis and transporting proteins into the nucleus. The shapes of the RNA level curves affected the estimation of gross transcription rates, and the crests and valleys of the RNA level curves responded to higher gross transcription rates. Different scenarios of sampling time intervals could change the shapes of the RNA level curves, resulting in different estimation values of gross transcription rates. Given the potential applications of the present method, further improvements are expected.

## 1. Introduction

Temporospatial transcription in organisms links genetic information with intra- and extraenvironmental factors, and transcriptional responses in biological and medical events exist in RNA abundance data (Ranz et al., 2003; Graveley et al., 2011; Kang et al., 2011; Hawrylycz et al., 2012; Kalucka et al., 2020; Armingol et al., 2021; Cho et al., 2021; Lee et al., 2021), which usually fluctuate because of the integration of the amount of transcription and RNA degradation (Wilusz et al., 2001; Schwanhäusser et al., 2011; Yamada and Akimitsu, 2019). RNA abundances directly detected by RT‒ qPCR, gene chip, RNA-seq, single-cell RNA-seq, etc., which is an RNA profile snapshot at a time point (La Manno et al., 2018; Marx, 2021), are often regarded as RNA amounts transcribed in the experimental period, thus possibly leading to incorrect results (Xu and Asakawa, 2019). To explore the molecular mechanisms of biological and medical phenomena, transcription rates should be obtained under the condition of RNA level fluctuations.

Special methods to directly examine transcription and RNA degradation rates have been developed in recent decades (Wada and Becskei, 2017; Wissink et al., 2019; Yamada and Akimitsu, 2019). For transcription rates, methods of transitioning from harmful to harmless labeled reagents, such as radioactive isotopes, 4-thiouridine (4sU), 5-ethynyluridine (EU), biotin, 5-bromo-uridine (BrU), and 2′-deoxy-2′-azidoguanosine (AzG), to mark nascent RNAs have been recently more broadly applied in transcriptome studies (Mukherjee and Beermann, 1965; Dölken et al., 2008; Jao and Salic, 2008; Tani et al., 2012; Kwak et al., 2013; Meng et al., 2020; Wan et al., 2021), but considering the ways in which RNA physically decays, e.g., some RNAs have half-lives of approximately 2 min or less (Baudrimont et al., 2017), the calculations can be difficult, in spite of the use of complicated operations to shorten sampling time, e.g., the commonly used shortest sampling time interval of 5 min (Schwalb et al., 2016), is one technique to address this problem. There are also nonlabeled methods to record transcripts by tracking bound RNA polymerases, blocking RNA-metabolizing enzymes, recording report marker signals, visualizing RNA dynamics, and so on (Gotta et al., 1991; Chubb et al., 2006; Singh and Padgett, 2009; Mayer et al., 2015; Shah et al., 2018; Rodermund et al., 2021). Similar methods of directly examining transcription rates are also used to directly detect RNA degradation rates (Bernstein et al., 2002; Yang et al., 2003; Deana et al., 2008; Rodermund et al., 2021). Unfortunately, the treatment of cells is a stimulus that impacts natural RNA level fluctuations, and these techniques are difficult to use in living organisms, especially humans, because of their disturbance of normal physiological processes.

Some computational methods to estimate transcription and RNA degradation rates have been recently reported for analyzing high-throughput gene expression data, including RNA-seq data. Mathematical models based on integrated data with enough intronic and exonic information have been developed to calculate the synthesis rates of premature RNA, the processing rates from premature RNA to mature RNA, and the degradation rates of mature RNA in eukaryotic organisms (Zeisel et al., 2011; Gray et al., 2014; Gaidatzis et al., 2015; Alkallas et al., 2017; Furlan et al., 2020). The models perform well in steady-state RNA level conditions. In the time course data of these studies, the RNA level data at different time points come from different cells because each sample or each cell in the high-throughput transcriptome experiments can only be detected one time, so the mature RNAs at a particular sampling time point are not the processing products from the premature RNAs at the previous sampling time points; thus, a derivation of premature RNAs at the previous sampling time point to mature RNAs at the next time point reduces the simulation ability and practical application of these methods, especially in the condition of RNA level fluctuations. These methods present the same problem when using different data from some independent studies to confirm the parameters of the derived relationship between RNA species. Moreover, methods that require high-throughput transcriptome data are not suitable for analyzing the data of common gene expression experiments, such as RT‒PCR. A method combining mathematical algorithms and experimental techniques to measure the metabolic rates of RNA in tissues seems reasonable (Schwalb et al., 2012; Blumberg et al., 2021), but the method cannot be used in transcriptomic detection in living organisms because they depend on labeled techniques, and it is difficult to estimate degradation rates in an unstable state. Calculation of transcription and RNA degradation rates with two time points are sometimes used (Huynh-Thu and Geurts, 2018), but the transcription and degradation events in the period between these two time points cannot be reasonably considered. In the time course data used in these computational methods, especially the RNA level fluctuation data, sampling time intervals are not considered. However, different sampling time intervals can affect the analysis results, e.g., changing the shapes of RNA level fluctuation curves (Xu and Askawa, 2021); if the sampling time interval is too large, the RNAs that are produced and then completely degraded between two adjacent sampling time points might be neglected in the analysis.

Previous studies might face the problem of the identification of complete nascent RNAs, which are the real direct products of transcription. RNAs enter the process of degradation following their production (Houseley and Tollervey, 2009). The amount of RNAs detected by the current techniques includes completely intact RNAs, partly edited RNAs, mature RNAs, microRNAs, and RNA degraded fragments, and RNA amounts detected in the experimental period include completely intact forms and incomplete forms of RNAs transcribed in the experimental period and the accumulated, incompletely degraded RNAs transcribed before the time of detection (La Manno et al., 2018; Xu and Askawa, 2019; Furlan et al., 2020). By using RNA abundance data detected by the current transcriptome tools, it is difficult to calculate the real transcription rates exactly. To address this problem, we defined the gross transcription rate in this study: RNA sequences detected during the experimental period are converted to complete nascent RNAs by their length, and the gross transcription rate equals the amount of all converted complete nascent RNAs divided by time. The gross transcription rate reflects the impact of transcriptional events before and in experiments on cells, for example, in the resource allocation of base and ribose. If the experimental data reflect the amounts of completely intact RNA, the gross transcription rates are the real transcription rates. In other cases, the gross transcription rate cannot completely represent the real transcription rate. The gross transcription rate has the significance of describing the role of the gene; for example, this parameter can indicate the cellular resource occupation of a gene in response to external stimuli. In fact, some units of RNA level in the results from current transcriptome tools, such as fluorescence signal intensity, RPKM (reads per kilobase per million reads), and TPM (transcripts per million), are analogs of converted complete nascent RNAs (Lipshutz et al., 1999; Mortazavi et al., 2008; Wagner et al., 2012). In this way, we can analyze various gene expression data by estimating gross transcription rates.

Therefore, we developed an algorithm to calculate gross transcription rates from RNA level fluctuation data obtained in various experimental conditions from routine gene expression assays to high-throughput transcriptome studies and to explore the less-studied problem of the effect of sampling time intervals on time course transcriptome experiments, aiming to provide a fast and practical computational tool for increasing gene expression analysis.

## Results

### 1.1 Effects of window length on the fitting between prediction and real data

In this algorithm, moving-window simulation was one of the main steps, and the setting of window length, which indicated how many values were used in a single simulation, could affect the results. Here, R-squared and median absolute percentage error (MdAPE) were the fitting indices to fit the prediction results to the real gene expression data. The results showed that in the analysis of expression data of gene YML021C (Figure 1a), as the window length increased from 3 to 17, the R-squared value generally decreased, the MdAPE value generally increased, and the simulation curves turned smoothly; when the window length was 3, the predictive curve fit the experimental data best, with the highest R-squared value of 0.985 and the lowest MdAPE value of 0.016.

**Figure 1.**
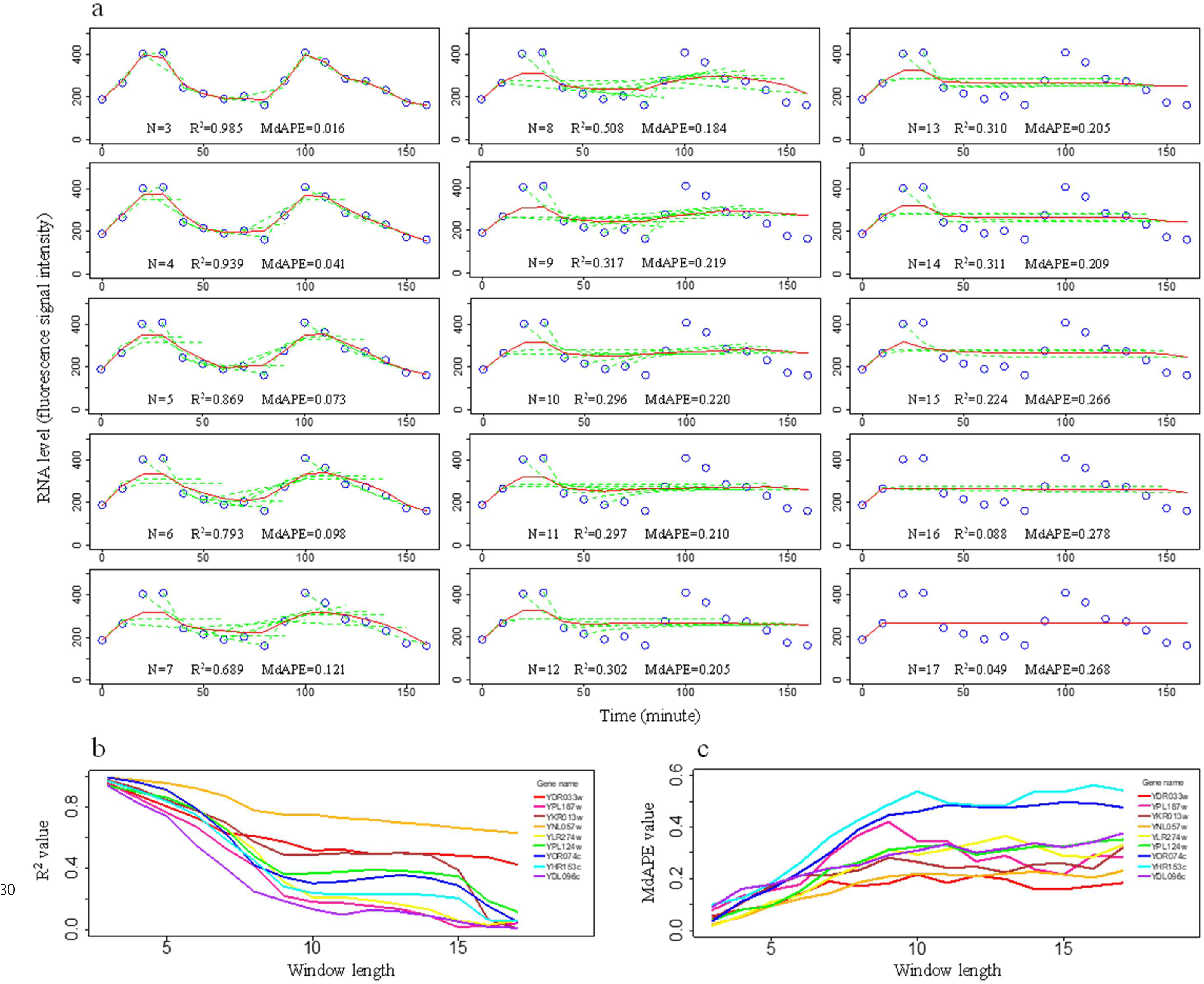
Effects of window length on the fitting between simulated and experimental values. The experimental data of the yeast RNA level were from the literature (Cho et al., 1998; Yueng et al., 2001). **a.** Effects of window length on the fitting between simulated and experimental values of the YML021C gene. Blue circles are experimental RNA levels of the yeast YML021C gene. The green dashed curves represent simulation curves for consecutive data points within a moving window. The red curves represent the mean RNA abundances of the points calculated in different moving windows. **b.** R-squared value as the index of fitting between simulated and experimental values of nine genes. The YDR033w gene had the first maximum gene expression mean of the experimental dataset of gene expression of 384 genes, the YPL187w gene had the second maximum gene expression mean, the YKR013w gene had the third maximum gene expression mean, the YDL096c gene had the first minimum gene expression mean, the YHR153c gene had the second minimum gene expression mean, the YOR074c gene had the third minimum gene expression mean, and the YNL057w, YLR274w, and YPL124w genes had the three median gene expression means. **c.** MdAPE value as the index of fitting between simulated and experimental values of nine genes. The nine genes are the same as in Figure 1b.

To study the effects of window length on analyzing an RNA level fluctuation dataset of 384 genes (Cho et al., 1998; Yeung et al., 2001), the first three maxima (YDR033w, YPL187w, and YKR013w), three medians (YNL057w, YLR274w, and YPL124w), and first three minima (YOR074c, YHR153c, and YDL096c) of gene expression means during the experimental period were selected to be representatives of the dataset. The results indicated that as the window length increased from 3 to 17, the R-squared value generally decreased (Figure 1b), and the MdAPE value generally increased (Figure 1c). In all nine gene expression datasets, when the window length was 3, the predictive curve fit the experimental data best, with the highest R-squared values and the lowest MdAPE values. Therefore, in the later analysis steps of this research, the window length was set to 3.

### 1.2 Estimation results of gross transcription and RNA degradation rates

Upon analyzing the dataset of 384 genes by the algorithm, the R-squared values were 0.859∼0.997, and the MdAPE values were 0.76%∼3.44% (Table S1). The simulation fit various fluctuated RNA abundance curves well (Figure 2a). These results indicated that gross transcription and RNA degradation rates could be calculated by using the present algorithm from RNA level fluctuation data. The absolute gross transcription rates were 19.5∼2645.2 fluorescence signal intensity/min, the absolute RNA degradation rates were 22.3∼2634.8 fluorescence signal intensity/min, and these absolute rates correlated with the means of RNA abundance over time (p<0.05). The relative gross transcription rates, which were normalized by dividing by the means, were 0.029∼2.268, and the relative RNA degradation rates were 0.039∼2.264. The absolute gross transcription rates were significantly correlated with the absolute RNA degradation rates (p<0.05), and the relative gross transcription rates were also significantly correlated with the relative RNA degradation rates (p<0.05).

**Figure 2.**
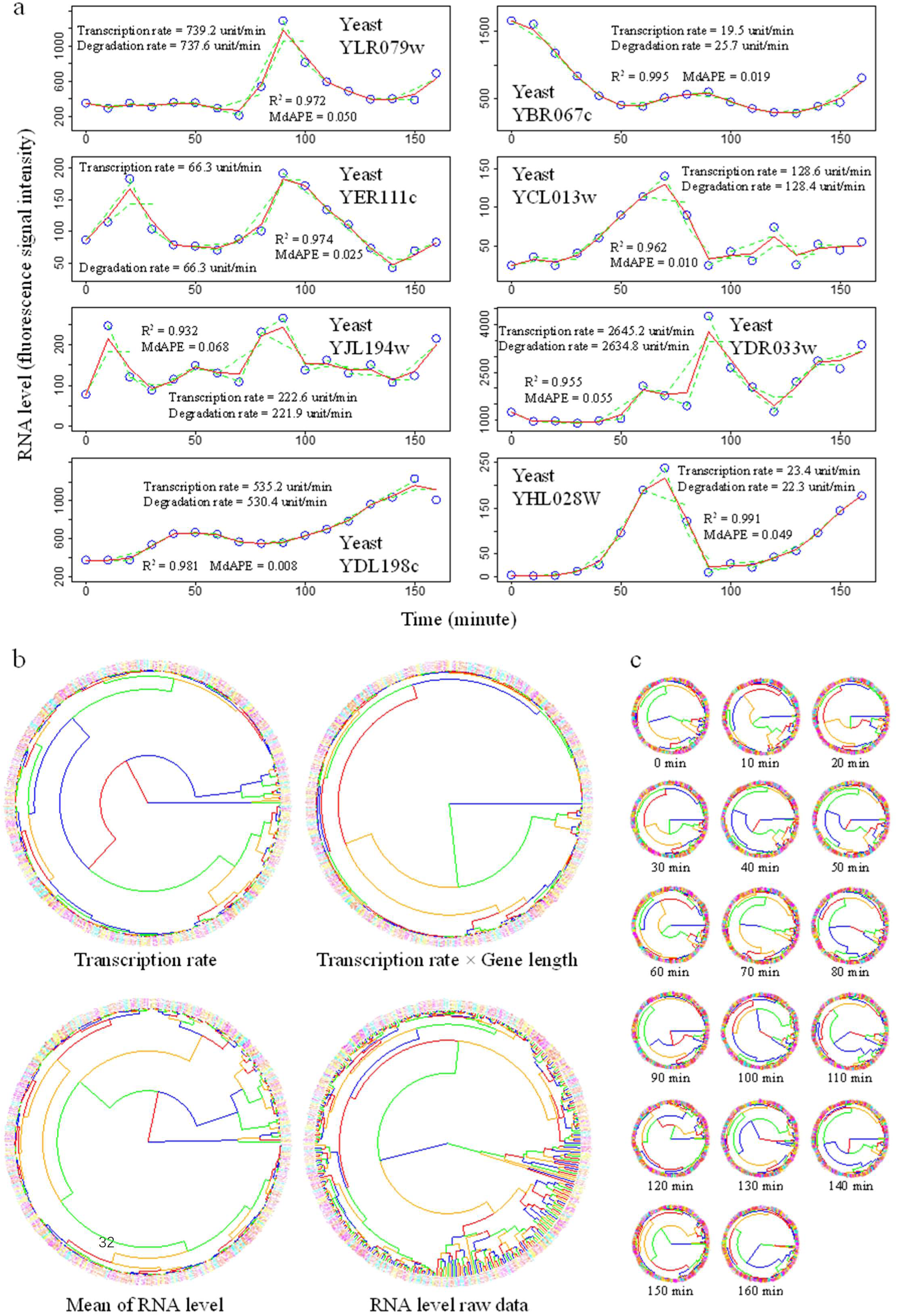
Estimation of gross transcription rates based on RNA level fluctuation data of yeast. The experimental data of the yeast RNA level were from the literature (Cho et al., 1998; Yueng et al., 2001). Transcription rates denoted gross transcription rates. **a.** Gross transcription rates and RNA degradation rates from several typical RNA level fluctuations. Blue circles are experimental data values. The green dashed curves represent simulation curves of a moving window with a window length of three. The red curves represent the mean RNA abundances of the points calculated in different moving windows. The unit was fluorescence signal intensity. The RNA level of the yeast gene YLR079w represented the one-crest curve. The RNA level of the yeast gene YER111c represented the two-crests curve. The RNA level of the yeast gene YJL194w represented the multiple-crests curve. The RNA level of the yeast gene YDL198c represented a smooth upward curve. The RNA level of yeast gene YBR067c represented a smooth downward curve, and it was the data with the minimum relative gross transcription rate, the minimum absolute gross transcription rate, and the minimum relative RNA degradation rate of the gene expression experimental dataset of 384 genes. The RNA level of the yeast gene YCL013w was the curve with the maximum relative gross transcription rate and the maximum relative RNA degradation rate. The RNA level of the yeast gene YDR033w was the curve with the maximum absolute gross transcription rate and the maximum absolute RNA degradation rate. The RNA level of the yeast gene YHL028W was the curve with the minimum absolute RNA degradation rate. **b.** Clustering analysis with the whole time-series RNA level dataset. **c.** Clustering analysis with the one-time-point RNA level data.

Cluster analysis results showed that the cluster structures based on gross transcription rates, gross transcription rates × gene length, RNA abundance fluctuations, and means of RNA abundance were different (Figure 2b). In the cluster result based on gross transcription rates, the 20 genes with the highest gross transcription rates (their locus tags were YDR033w, YPL187w, YBL003c, YAL040c, YKL066W, YBL002w, YNR016c, YLR286c, YLR297W, YKR013w, YBR073w, YJL173c, YJR148w, YMR256c, YLR254c, YLR050c, YPR120c, YBR158w, YDR309c, and YMR199w, with gene symbols of MRH1, MF(ALPHA)1, HTA2, CLN3, YKL066W, HTB2, ACC1, CTS1, YLR297W, PRY2, RDH54, RFA3, BAT2, COX7, NDL1, EMA19, CLB5, AMN1, GIC2, and CLN1, respectively) were grouped in an integrated cluster; their functions were mainly related to activities of ion channel, mating pheromone, DNA binding, cyclin-dependent protein regulator, carboxylase, endochitinase, sterol binding, transaminase, oxidase, microtubule binding, regulation of protein targeting to mitochondrion, and GTPase binding, which play roles in cell division regulation, DNA replication, biosynthesis of protein and fatting acid, material transport, energy metabolism, and cell fusion. The 8 genes listed in the top 10 gross transcription rates (YDR033w, YPL187w, YBL003c, YAL040c, YBL002w, YNR016c, YLR297W, YKR013w) are listed in the top 10 means of RNA level (Table S1); the functions of YKL066W and YLR297W seemed unclear. In the cluster result based on relative gross transcription rates, the 5 genes with the highest relative gross transcription rates (their locus tags were YCL013w, YKL066W, YLR233c, YLR286c, and YDL181w, with gene symbols of YCL013w, YKL066W, EST1, CTS1, and INH1, respectively) were grouped in an integrated cluster; their functions were mainly related to activities of DNA binding, endochitinase, and ATPase inhibitor, which play roles in cell fusion, DNA replication, and energy metabolism; the functions of YKL066W seemed unclear, and YCL013w was currently a deleted ORF. In the cluster result based on gross transcription rate × gene length (related to the consumption of cellular resources, such as base, ribose, and energy), the highest gross transcription rate × gene length grouped in a single cluster was YNR016c (its gene symbol was ACC1), and its function was fatty acid biosynthesis and transporting protein into the nucleus. In the cluster result based on the mean of fluctuated RNA abundance, the 6 genes with the highest means (their locus tags were YDR033w, YPL187w, YKR013w, YBL003c, YBL002w, and YNR016c, with gene symbols of MRH1, MF(ALPHA)1, PRY2, HTA2, HTB2, and ACC1, respectively) were grouped in an integrated cluster; their functions were mainly related to activities of ion channel, mating pheromone, sterol binding, DNA binding, and carboxylase, which play roles in material transport, cell fusion, DNA replication, and biosynthesis of fatting acid. The cluster results based on raw data of RNA abundance seemed to have more detailed branches; there were 19 genes in the list of the 24 highest means of fluctuated RNA abundance (their locus tags were YDR033w, YPL187w, YKR013w, YBL003c, YBL002w, YNR016c, YLR297W, YAL040c, YLR050c, YJL173c, YGR189C, YBR073w, YBR088c, YPL256c, YMR003w, YMR199w, YMR215w, YJR076c, and YBR071w) in an integrated cluster.

RNA level data at each sampling time point could also be used to perform clustering analysis. The results showed that the cluster results at each time point were different (Figure 2c), and they were also different from the results analyzed with the data from all time points (Figure 2b). This indicated that the experimental data of the RNA level snapshot at one time point made it difficult to extrapolate the results over the entire experimental period.

### 1.3 Effects of the shapes of RNA level curves on the estimations of the gross transcription rates

The shapes of RNA level fluctuations affected the estimation values of the gross transcription rates (Figure 3). The gross transcription rates of the RNA level curves with multiple crests or valleys seemed to be higher than those with fewer crests or valleys. In the crests and valleys of the three window length scenario, there were higher first transcription coefficients (the transcription rate coefficients in the first moving windows) or second transcription coefficients (the transcription rate coefficients in the second moving windows), accompanied by higher first RNA degradation rate coefficients or second RNA degradation rate coefficients, and the crests and valleys of the RNA level curves corresponded to the rapidly rising parts in the curves of net RNA production accumulation (Figure 3a). Because the estimation value of a time point was the mean of the estimation values of the same point under different overlap moving windows, a high estimation value from any moving window, caused by the first transcription rate coefficients or the second transcription rate coefficients, would lead to a higher mean value.

**Figure 3.**
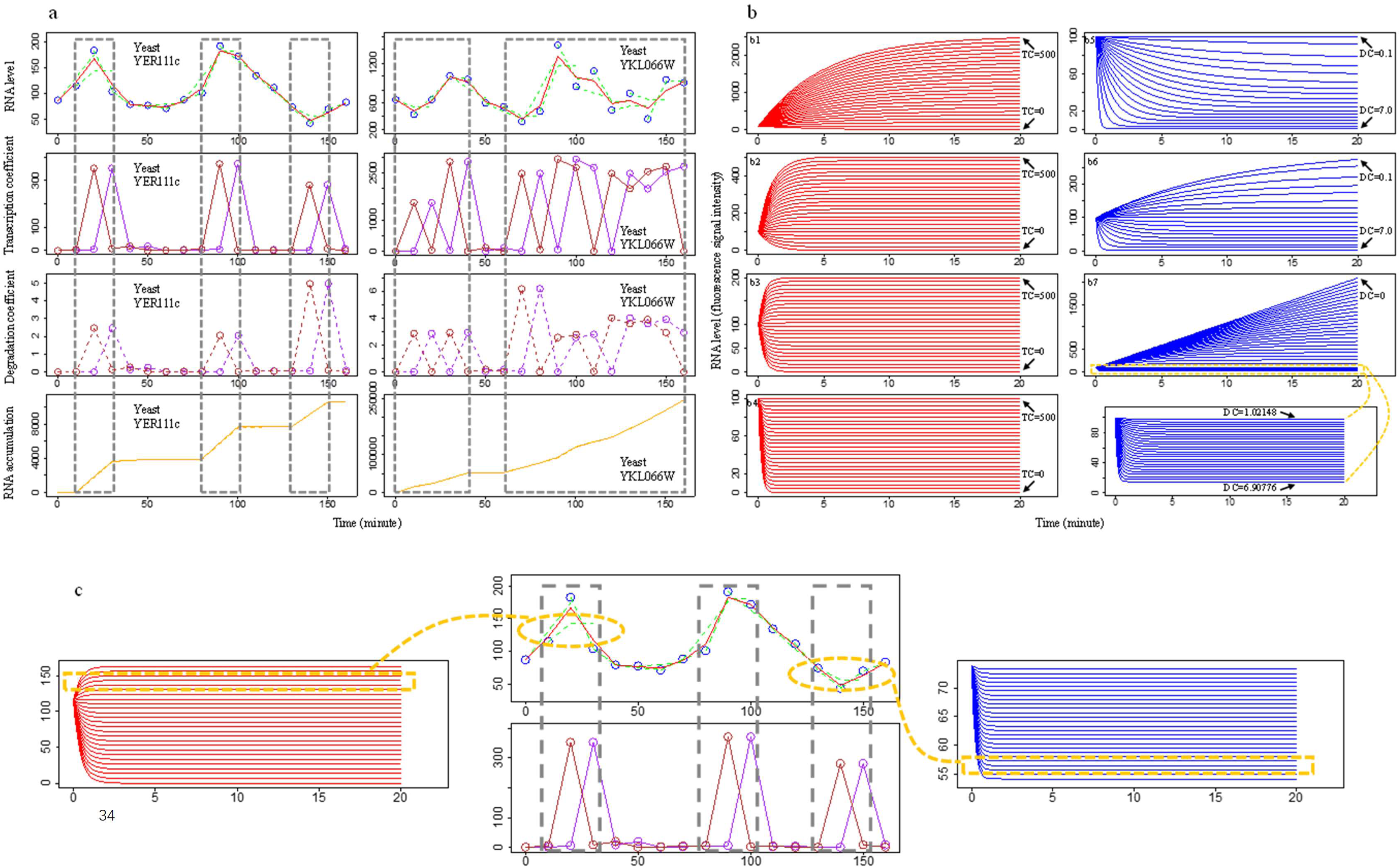
Effects of the shapes of RNA level curves on the estimations of gross transcription rates. The experimental data of the yeast RNA level were from the literature (Cho et al., 1998; Yueng et al., 2001). Transcription rates denoted gross transcription rates. **a.** Effects of crests and valleys of RNA levels on the estimation of gross transcription rate. Blue circles are experimental RNA levels of the yeast YER111c gene and YEL066W gene. The green dashed curves represent simulation curves of a moving window with a window length of three. The red curves represent the mean RNA abundances of the points calculated in different moving windows. The solid purple curves represent the transcription coefficients in the first moving window, and the solid brown curves represent the transcription coefficients in the second moving window. The dashed purple curves represent the degradation coefficients in the first moving window, and the dashed brown curves represent the degradation coefficients in the second moving window. The solid orange curves represent transcription accumulation, and the dashed orange curves represent degradation accumulation. The unit of the y-axis of the RNA level is fluorescence signal intensity. The unit of the y-axis of the transcription coefficient is fluorescence signal intensity/min. The unit of the y-axis of transcription accumulation is fluorescence signal intensity. **b.** Shapes of the model curves with different coefficients of transcription and RNA degradation. If the RNA degradation coefficient=0, then the model was Equation 5 in the Methods section. If the RNA degradation coefficient≠0, then the model was Equation 4 in the Methods section. Red curves are the curves of the model with fixed values for RNA degradation coefficients and changed values for transcription coefficients. Blue curves are the curves of the model with fixed values for transcription coefficients and changed values for RNA degradation coefficients. TC denotes the transcription coefficient. DC denotes the RNA degradation coefficient. **b1**, RNA degradation coefficient=0.2, transcription coefficient=0∼500, N_0_=100. The transcription coefficients of the model curves from down to top were 0, 20, 40, 60, 80, 100, 120, 140, 160, 180, 200, 220, 240, 260, 280, 300, 320, 340, 360, 380, 400, 420, 440, 460, 480, and 500. **b2**, RNA degradation coefficient=1.0, transcription coefficient=0∼500, N_0_=100. The transcription coefficients of the model curves from down to top were 0, 20, 40, 60, 80, 100, 120, 140, 160, 180, 200, 220, 240, 260, 280, 300, 320, 340, 360, 380, 400, 420, 440, 460, 480, and 500. **b3**, RNA degradation coefficient=2.5, transcription coefficient=0∼500, N_0_=100. The transcription coefficients of the model curves from down to top were 0, 20, 40, 60, 80, 100, 120, 140, 160, 180, 200, 220, 240, 260, 280, 300, 320, 340, 360, 380, 400, 420, 440, 460, 480, and 500. **b4**, RNA degradation coefficient=5.0, transcription coefficient=0∼500, N_0_=100. The transcription coefficients of the model curves from down to top were 0, 20, 40, 60, 80, 100, 120, 140, 160, 180, 200, 220, 240, 260, 280, 300, 320, 340, 360, 380, 400, 420, 440, 460, 480, and 500. **b5**, Transcription coefficient=10. RNA degradation coefficient=0.1∼7.0, N_0_=100. The RNA degradation coefficients of the model curves from top to bottom are 0.10, 0.11, 0.13, 0.15, 0.17, 0.20, 0.23, 0.26, 0.30, 0.35, 0.41, 0.50, 0.60, 0.75, 1.0, 1.5, 2.5, and 7.0. **b6**, Transcription coefficient=30. RNA degradation coefficient=0.1∼7.0, N_0_=100. The RNA degradation coefficients of the model curves from top to bottom are 0.10, 0.11, 0.13, 0.15, 0.17, 0.20, 0.23, 0.26, 0.30, 0.35, 0.41, 0.50, 0.60, 0.75, 1.0, 1.5, 2.5, and 7.0. **b7**, Transcription coefficient=100. RNA degradation coefficient=0∼6.90776, N_0_=100. The RNA degradation coefficients of the model curves from top to bottom are 0, 0.00389597, 0.008013, 0.0123787, 0.0170207, 0.0219736, 0.0272718, 0.0329776, 0.0391462, 0.0458468, 0.0531828, 0.061258, 0.0702242, 0.0802957, 0.0917212, 0.10488, 0.120312, 0.138791, 0.161572, 0.190709, 0.229911, 0.286561, 0.377606, 0.551515, 1.02148, 1.05753, 1.09621, 1.13783, 1.18273, 1.23133, 1.28409, 1.34158, 1.40445, 1.4735, 1.54969, 1.6342, 1.72845, 1.83425, 1.95383, 2.09009, 2.24679, 2.42889, 2.64311, 2.89877, 3.20918, 3.59405, 4.0838, 4.7281, 5.61378, and 6.90776. **c.** Model curves with convexity and concavity simulating the crests and valleys of the RNA levels. The plots in the middle are the experimental RNA levels of the yeast YER111c gene, with the results of the model simulations and transcription coefficient analysis. The red plot on the left shows the model curves with a fixed value of the RNA degradation coefficient and changed values of the transcription coefficients (according to the crest circled with a dashed line, the RNA degradation coefficient was 2.45225, and the transcription coefficients were 0∼400). The blue plot on the right shows the curves of the model with a fixed value of the transcription coefficient and changed values of the RNA degradation coefficients (according to the valley circled with a dashed line, the transcription coefficient was 280.617, and the RNA degradation coefficients were 3.827∼5.177).

The results under the simulation background may help to explain the effects of the shapes of RNA level curves on the estimation of gross transcription and degradation rates. When the degradation coefficients were stable, the transcription coefficients became lower as the shapes of the RNA level curves decreased from convexly upward to concavely downward (Figure 3b). When the transcription coefficients were stable, the degradation coefficients increased as the shapes of the RNA level curves increased from convexly upward to concavely downward.

In the crests of the RNA level, the predicted RNA level curves seemed to respond to the convexly upward parts, and there were both high transcription rates and high degradation rates. In terms of biological significance, this could be considered because the RNA amounts underwent high transcription rates, immediately followed by high RNA degradation rates (Figure 3c). In the points of valleys, the predicted RNA level curves seemed to respond to the concavely downward parts, and there were both high transcription rates and high degradation rates, but the biological significance could be considered as the high RNA degradation rates first, immediately followed by high transcription rates.

### 1.4 Effects of sampling time intervals on the estimations of the gross transcription rates

Sampling time intervals played an important role in the estimations of gross transcription rates, and different scenarios of sampling time intervals resulted in different estimation values of gross transcription rates (Figure 4). There were similar effects of sampling time intervals on RNA degradation rates.

**Figure 4.**
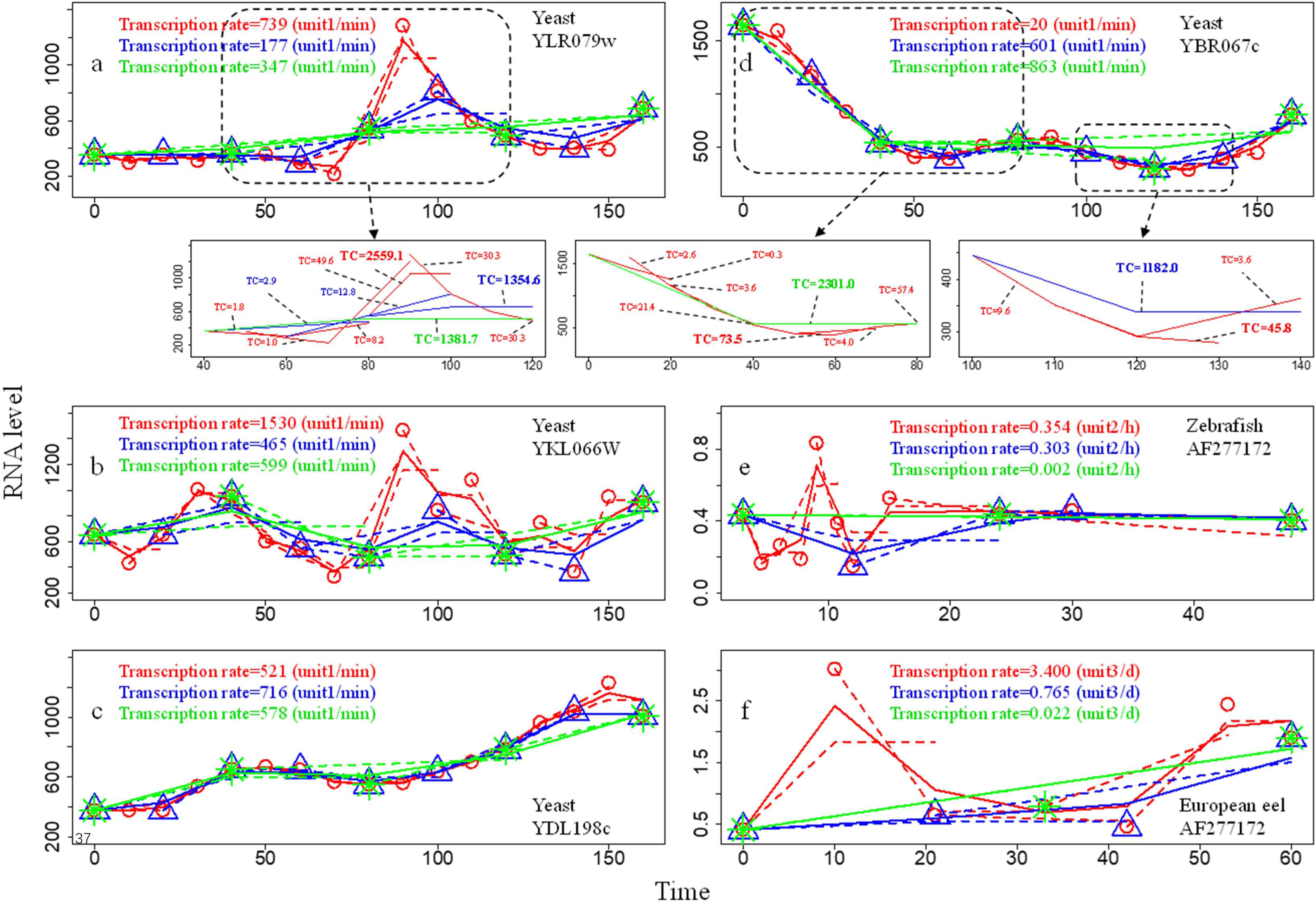
Effect of sampling time intervals on the estimations of gross transcription rates. In a, b, c, and d, the results of data with a sampling time interval of 10 minutes are shown in red, the results of data with a sampling time interval of 20 minutes are shown in blue, and the results of data with a sampling time interval of 40 minutes are shown in green. In e, the results of data with sampling time intervals of 1.3-18 hours are shown in red, the results of data with sampling time intervals of 6-18 hours are shown in blue, and the results of data with sampling time intervals of 21-24 hours are shown in green. In f, the results of data with sampling time intervals of 7-12 days are shown in red, the results of data with sampling time intervals of 18-21 days are shown in blue, and the results of data with sampling time intervals of 27-33 days are shown in green. The circles, triangles, and asterisks are experimental data points with different sampling time intervals. The dashed curves represent simulation curves of a moving window with a window length of three. Unit 1 was fluorescence signal intensity, unit 2 was normalization of fluorescence signal intensity, and unit 3 was relative quantification of quantitative RT‒PCR. Transcription rates denoted gross transcription rates. **a.** One-crest RNA level fluctuation data: effects of sampling time intervals on the estimations of gross transcription rates of the yeast YLR079w gene. The experimental data were from the literature (Cho et al., 1998; Yeung et al., 2001). The solid curves in the large plot represent the mean RNA abundances of the points calculated in different moving windows. The solid curves in the small plot represent simulation curves for three consecutive points. The x-axis is time, and the unit of time is minutes. The y-axis is the RNA level, and the unit is the fluorescence signal intensity. **b.** Multiple-crests RNA level fluctuation data: effects of sampling time intervals on the estimations of gross transcription rates of the yeast YKL066W gene. The experimental data were from the literature (Cho et al., 1998; Yeung et al., 2001). The solid curves represent the mean RNA abundances of the points calculated in different moving windows. The x-axis is time, and the unit of time is minutes. The y-axis is the RNA level, and the unit is the fluorescence signal intensity. **c.** Smooth upward RNA level fluctuation data: effects of sampling time intervals on the estimations of gross transcription rates of the yeast YDL198c gene. The experimental data were from the literature (Cho et al., 1998; Yeung et al., 2001). The solid curves represent the mean RNA abundances of the points calculated in different moving windows. The x-axis is time, and the unit of time is minutes. The y-axis is the RNA level, and the unit is the fluorescence signal intensity. **d.** Smooth downward RNA level fluctuation data: effects of sampling time intervals on the estimations of gross transcription rates of the yeast YBR067c gene. The experimental data were from the literature (Cho et al., 1998; Yeung et al., 2001). The solid curves in the large plot show the mean RNA abundances of the points calculated in different moving windows. The solid curves in the small plots represent simulation curves for three consecutive values. The x-axis is time, and the unit of time is minutes. The y-axis is the RNA level, and the unit is the fluorescence signal intensity. **e.** Effects of sampling time intervals on the estimations of gross transcription rates of the zebrafish AF277172 gene. The experimental data were from the literature (Mathavan et al., 2005). The solid curves represent the mean RNA abundances of the points calculated in different moving windows. The x-axis is time, and the unit of time is hours. The y-axis is the RNA level, and the unit is normalization of the fluorescence signal intensity. **f.** Effects of sampling time intervals on the estimations of gross transcription rates of the European eel Cathf gene. The experimental data were from the literature (Bolliet et al., 2017). The solid curves represent the mean RNA abundances of the points calculated in different moving windows. The x-axis is time, and the unit of time is days. The y-axis is the RNA level, and the unit is the relative quantification of quantitative RT‒PCR.

Different sampling time intervals changed the shapes of the RNA level curves. In the curves with crests and valleys, longer sampling time intervals could make the crest of the curve smoother (Figure 4a, Figure 4b, Figure 4e, and Figure 4f). The effects of sampling time intervals on the estimation of gross transcription rates could be complicated. Taking 9 time points of the RNA level curve of yeast gene YLR079w as an example (the small plot of Figure 4a), with the sampling interval of 10 minutes in this crest, there was only one high transcription coefficient over 2500, and the other six transcription coefficients were less than 50, resulting in the lack of a high mean of the transcription coefficient within this part; with the sampling interval of 20 minutes, there was one high transcription coefficient over 1000, and the other two transcription coefficients were less than 20; with the sampling interval of 40 minutes, there was only one transcription coefficient whose value was over 1000; so the highest mean transcription coefficient in this part was the one with a sampling time interval of 40 minutes, and the lowest one was that with a sampling time interval of 20 minutes. In the relatively smooth curves, longer sampling time intervals could sometimes result in higher transcription coefficients (Figure 4c and Figure 4d), partly because the look-like smooth curves actually had smooth crests and valleys, and longer sampling time intervals might make crests and valleys longer. In the first 80 minutes of a smoothly downward curve of the yeast gene YBR067c RNA level (the small plot on the left of Figure 4d), if the sampling time interval was 10 minutes, the transcription coefficients were less than 100, resulting in a low mean transcription coefficient during this experimental period; however, if the sampling time interval was 40 minutes, the RNA amounts at the 0^th^, 40^th^, and 80^th^ minutes were 1649, 540, and 563 fluorescence signal intensities, respectively, forming a valley. Thus, a simulation curve with a larger degree of concavely downward depression was formed, leading to a transcription coefficient over 2000 during this experimental period. From the 100^th^ to the 140^th^ minute of a smoothly downward curve of the yeast gene YBR067c RNA level (the small plot on the right of Figure 4d), the transcription coefficients were less than 100 if the sampling time interval was 10 minutes; however, if the sampling time interval was 20 minutes, the RNA amounts at the 100^th^, 120^th^, and 140^th^ minutes were 446, 291, and 384 fluorescence signal intensities, respectively, forming a valley and leading to a transcription coefficient over 1000 during this experimental period. Therefore, the effects of the sampling time interval might change in different parts of a curve, and the effects of the sampling time interval on the whole curve should integrate each effect on different parts of the curve (Figure 4d).

If the sampling time interval was too long, the simulation curves would miss the details of crests and valleys of the RNA levels, resulting in different gross transcription rates. In the RNA level data of the zebrafish AF277172 gene, the gross transcription rate with a sampling time interval of 21-24 hours was less than 1% of that with a sampling time interval of 1.3-18 hours (Figure 4e). In the RNA level data of the European eel Cathf gene, although some values of gross transcription rates with different sampling time interval designs were calculated, it seemed that the algorithm in the present study could not analyze these data because the sampling time interval of the original experimental data was too long — at least one week (Figure 4f). The RNA half-lives of eukaryotes may be less than 2 days (McManus et al., 2015); therefore, many undetectable RNAs may be produced and completely degraded within one week, resulting in underestimated gross transcription rates.

### 1.5 Effects of the number of sampling time points on the simulation fitting and the gross transcription rate estimation

The number of sampling time points would affect the results of the simulation and related analysis. When the number of sampling time points of yeast YJL194w gene expression data ranged from 3 to 17, the ranges of gross transcription rates, R-squared values, and MdAPE values were 172.0-488.0 fluorescence signal intensity/min, 0.487-0.951, and 0.028-0.254, respectively (Figure 5a). Adding certain sampling time points, such as the fourth and sixth sampling time points in yeast YJL194w gene expression data, could make the relative parameters change significantly. Too small a number of sampling time points sometimes might lead to a simulated curve with a lower fit; for example, when the number of sampling time points in the yeast YJL194w gene expression data was 3, the simulated values did not fit the experimental data well, with an R-squared value of 0.487 and an MdAPE value of 0.254.

**Figure 5.**
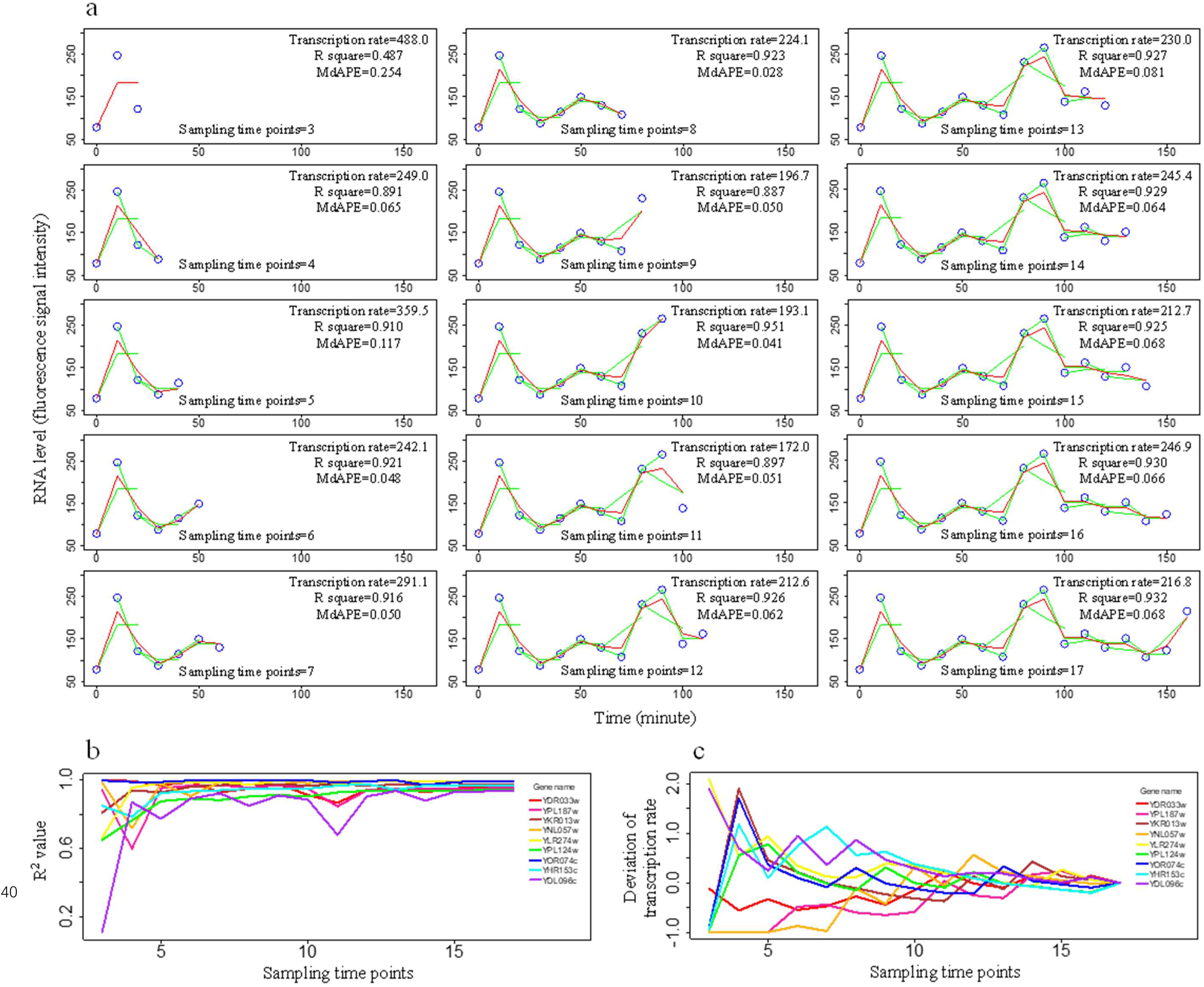
Effects of the number of sampling time points on the estimations of gross transcription rates and the fitting indices. The experimental data of the yeast RNA level were from the literature (Cho et al., 1998; Yueng et al., 2001). Transcription rates denote gross transcription rates. **a.** Estimation of gross transcription rates by using different sampling time points. Blue circles are experimental RNA levels of the yeast YJL194w gene. The green curves represent simulation curves of a moving window with a window length of three. The red curves represent the mean RNA abundances of the points calculated in different moving windows. The unit of gross transcription rate is fluorescence signal intensity/min. **b.** R-squared values as the index of fitting between simulated and experimental values of nine genes. The YDR033w gene had the first maximum gene expression mean of the experimental dataset of gene expression of 384 genes, the YPL187w gene had the second maximum gene expression mean, the YKR013w gene had the third maximum gene expression mean, the YDL096c gene had the first minimum gene expression mean, the YHR153c gene had the second minimum gene expression mean, the YOR074c gene had the third minimum gene expression mean, and the YNL057w, YLR274w, and YPL124w genes had the three median gene expression means. **c.** Deviation of gross transcription rates for the nine genes by using different sampling time points. Deviation of gross transcription rate = (Gross transcription rate by using certain sampling time points – Gross transcription rate by using the maximum sampling time points)/Gross transcription rate by using the maximum sampling time points. The nine genes are the same as in Figure 5b.

To study the effects of the number of sampling time points on the simulation fitting and the gross transcription rate estimation with an RNA level fluctuation dataset of 384 genes (Cho et al., 1998; Yeung et al., 2001), the first three maxima, three medians, and the first three minima of gene expression means during the experimental period were selected to be representatives of the dataset (Figure 5b and Figure 5c). Generally, R-squared values seemed to increase with the addition of sampling time points in order from 3 to 17 (Figure 5b). The changes in R-squared values might fluctuate depending on the RNA level fluctuations, such as adding the eleventh and fourteenth sampling time points of the gene YDL096c; however, with the increase in the sampling time point number, the effects of certain data points on R-squared values were diluted. Adding new sampling time points could change the estimated gross transcription rates (Figure 5c). The differences between the estimation of transcription rate with 3 sampling time points and that with 17 sampling time points were large, with some deviation of gross transcription rates reaching 200%; by adding sampling time points, the deviations of gross transcription rates between a certain time point and the end of the experiment became progressively smaller. Similarly, certain individual sampling time points could affect the estimations of gross transcription rates, but these effects were diluted by the increase in the sampling time point number.

### 1.6 Effects of RNA level fluctuations under a simulation background

The algorithm could simulate smooth curves, such as a straight line function curve, exponential function curve, fractional function curve, parabolic function curve, and sine function curve, with a good fitting index of high R-squared values greater than 0.99 and low MdAPE values less than 0.03 (Figure 6a, Figure 6b, Figure 6c, Figure 6d, and Figure 6e). Compared to the straight line function curve, the other four smooth curves, which had larger curvatures or had crests and valleys, did not affect the good fitting index but led to higher relative gross transcription rates; the relative gross transcription rates of the exponential function curve, the fractional function curve, the parabolic function curve, and sine function curve were 0.04, 0.03, 0.02, and 0.32, respectively, but that of the straight line function curve was only 0.01.

**Figure 6.**
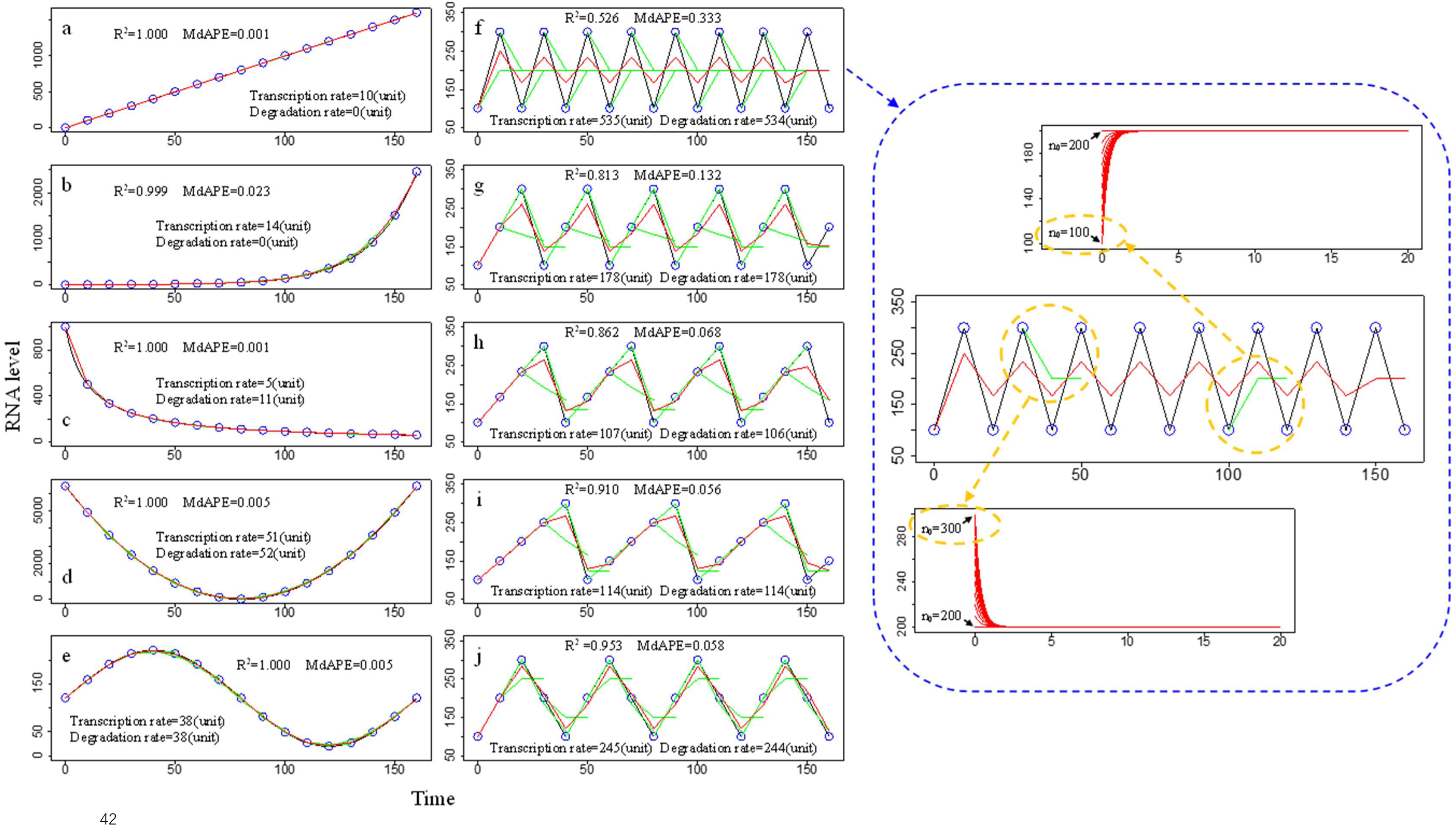
Simulation of data points from different function curves. The black curves represent the function curves, and blue circles are data points on the curve. The green curves represent simulation curves of a moving window with a window length of three. The red curves represent the mean RNA abundances of the points calculated in different moving windows. The data were simulation background data without specific units of time, RNA level, gross transcription rate, and degradation rate. Transcription rates denoted gross transcription rates. **a.** Simulation of data points on the curve of y=10x. **b.** Simulation of data points on the curve of y=1.05^x^. **c.** Simulation of data points on the curve of y=10000/(10+x). **d.** Simulation of data points on the curve of y=(x-80)^2^. **e.** Simulation of data points on the curve of y=100sin(0.039269908x)+120 **f.** Simulation of data points on the curve of a periodic function with two data points as a cycle, and the order of the data points in the cycle is 100 and 300. In the simulation curves in red in the right top plot, the transcription coefficient was 534.66, the degradation coefficient was 2.6733, and the values of N_0_ from bottom to top were 100, 110, 120, 130, 140, 150, 160, 170, 180, 190, and 200; except that when N_0_=200, the curve was a horizontal line, and all others were curves of monotonically increasing functions. In the simulation curves in red in the bottom right plot, the transcription coefficient value was 534.66, the degradation coefficient value was 2.6733, and the values of N_0_ from bottom to top were 200, 210, 220, 230, 240, 250, 260, 270, 280, 290, and 300; except that when N_0_=200, the curve was a horizontal line, and all others were curves of monotonically decreasing functions. **g.** Simulation of data points on the curve of a periodic function with three data points as a cycle, and the order of the data points in the cycle is 100, 200, and 300. **h.** Simulation of data points on the curve of a periodic function with four data points as a cycle, and the order of the data points in the cycle is 100, 167, 233, and 300. **i.** Simulation of data points on the curve of a periodic function with four data points as a cycle, and the order of the data points in the cycle is 100, 150, 200, 250, and 300. **j.** Simulation of data points on the curve of a periodic function with four data points as a cycle, and the order of the data points in the cycle is 100, 200, 300, and 200.

Every 3 consecutive sampling time points were simulated by a basic equation of the present algorithm, which was a monotonic function, either monotonically increasing (the right top plot of Figure 6f) or monotonically decreasing (the right bottom plot of Figure 6f); thus, if these 3 sampling time points formed a crest or a valley, the monotonic equation did not fit such a fluctuating curve well. Every three consecutive points of the curve in Figure 6f formed a crest or a valley, so the algorithm could not simulate the data well, with a low R-squared value and a high MdAPE value. If a crest or a valley was formed by four consecutive values (Figure 6g), five consecutive values (Figure 6h and Figure 6j), or six consecutive values (Figure 6i), this algorithm could fit the data better, with higher R-squared values and lower MdAPE values, partly because the predicted results from 3 consecutive points that did not form a crest or a valley diluted the results from 3 consecutive points that formed a crest or a valley. The more evenly distributed the points on both sides of the crest or valley, the better fitting index was obtained; thus, the fitting index of Figure 6j was better than that of Figure 6h. The presence of crests and valleys in the curves increased transcription and degradation coefficients in the models, so the relative gross transcription rates of Figure 6f, Figure 6g, Figure 6h, Figure 6i, and Figure 6j were 2.75, 0.92, 0.55, 0.60, and 1.26, respectively, higher than those of the smooth curves (Figure 6a, Figure 6b, Figure 6c, Figure 6d, and Figure 6e), which were less than 0.32.

## 2 Discussion

### 2.1 Calculation of gross transcription rates from RNA level fluctuation data

Transcriptomic datasets are accumulatively large, and some of them contain consecutive sampling data with various RNA level fluctuations, which enables the application of the present algorithm to calculate gross transcription rates and RNA degradation rates in most cases. The calculation of gross transcription rates and RNA degradation rates of a published dataset could help to discover other new findings (Figure 2). The parameter of gross transcription rate × gene length could represent the consumption of cellular resources for gene expression during the experimental period, reflecting the investment and resource allocation of a gene product in cells. Gene YNR016c was found to be the maximum of gross transcription rate × gene length, approximately three times the second maximum gene. The main function of the YNR016c gene is fatty acid biosynthesis and the control of lipogenic enzymes (Roggenkamp et al., 1980; Galdieri and Vancura, 2012; Blank et al., 2017). Why did cells allocate maximum resources to the YNR016c gene? Was it because the formation of cell membranes and intracellular membrane structures requires a large amount of fatty acids? Obviously, this was an interesting question that deserves further study. Absolute gross transcription rates were generally closely related to the means of RNA level (P<0.05), and relative gross transcription rates seemed to be based on fluctuations of RNA levels, such as the numbers and the slopes of crests and valleys in RNA level curves. The 10 maximum absolute gross transcription rates in the dataset of 384 genes were generally listed in the top 10 means of RNA level, except for gene YKL066W and gene YLR286c. These two genes were listed at order 43 and order 47 in the means of the RNA level, but they were listed at order 2 and order 4 in the relative gross transcription rates, suggesting that their higher lists in the absolute gross transcription rates mainly resulted from the larger fluctuations of RNA level curves. Why gene YKL066W, which seemed unlikely to encode a functional protein (Fisk et al., 2006; Stuart et al., 2009), had the high gross transcription rates needs further study.

The analysis of temporal patterns of gene expression from RNA level fluctuations was based on amounts of RNA transcribed in a certain time period, but time series RNA snapshots were usually used to assess temporal gene expression patterning (DeRisi et al., 1997; Kasturi et al., 2003; Bar-Joseph et al., 2012), partly resulting from the limited data of continuous transcription rates available. Time series gene expression profiles were mixed with temporal patterns of RNA degradation. The present algorithm could calculate temporal gene expression patterns in different time ranges from RNA level fluctuation data (Figure S1). Transcription and RNA degradation accumulation curves (Figure S1b) could provide more useful information than temporal gene expression, so the accumulation patterns of transcription and RNA degradation are worthy of further study. Only gross transcription rates through the experimental periods, accompanied by RNA degradation rates, could sometimes describe gene expression trends over time. For relatively smooth RNA level curves, the upward curves might be explained as gross transcription rates higher than RNA degradation rates (Figure 2a gene YDL198c, Figure 6a, Figure 6b), and the downward curves might be explained as transcription rates lower than RNA degradation rates (Figure 2a gene YBR067c, Figure 6c). Relative gross transcription rates could be used to compare fluctuations of RNA level curves, and higher relative gross transcription rates usually meant larger fluctuations. If the means of the RNA levels were similar, absolute gross transcription rates could be used to compare fluctuations in the RNA level curves (Figure S1). Gross transcription rates can also be used to distinguish differences in time series gene expression experiments. In some time-series gene expression experiments, if the gene expression curves of different treatment groups are interleaved, such as the curves with similar means in Figure S1a, the statistical method might judge that they were not significantly different, but gross transcription rates calculated from the data might have the potential to differentiate the effects of different treatments according to the detailed shapes of RNA level curves (Figure S1b, Figure S1c, Figure S1d, and Figure S1e).

The simulation equation regarding transcription (Equation 1 in the Methods section) was based on the assumption that mRNA is produced at a constant rate (Palumbo et al., 2015; Yamada and Akimitsu, 2019), and then the transcription coefficient should be stable. This assumption is statistical or theoretical, but whether it can be applied to a specific case remains unknown. In a cell, transcription at a constant rate within a certain time can be explained by the RNA polymerase lining up on the same DNA template at the same interval for the transcription reaction, but cellular transcription might be stochastic and pulsing partly due to the random collisions and reactions of biomolecules such as DNA, RNA polymerase or other factors (Kærn et al., 2005; Chubb et al., 2006; Locke et al., 2011). Thus, whether mRNAs of cells in tissues are produced at a constant rate under specific conditions needs to be proven by more data.

This algorithm has prospects for application. The basic models of this algorithm (Equation 4 and Equation 5 in the Methods section) could only theoretically explain the changes in RNA abundance over a short time, so the simulation curves might be dramatically different from the experimental data of RNA level fluctuations over a longer time; the reason was that the transcription and RNA degradation coefficients were constants within a short time, but there were different coefficients of transcription and RNA degradation in the case of RNA level fluctuation data. The design of the moving window in the present algorithm resolved this problem. The present method can utilize time series gene expression data from most methods, such as RT‒qPCR, gene chip, and RNA-seq. By using an exhaustive search method to select appropriate transcription and RNA degradation coefficients, the time complexity of this algorithm was O(n^4^); for data of normal gene expression experiments within 100 continuous sampling time points, the computer operation speed of this algorithm is relatively fast.

### 2.2 Effects of the shapes of RNA level curves

The curves of RNA level fluctuations might have various shapes, and through the moving windows, this algorithm could take local sections of RNA level curves as the analysis units to perform simulations, thus making the predicted results fit the real data better (Figure 1). The shape of the RNA level curve section for direct simulation was a major impactor of the fitting between simulation curves and experimental curves. Each local curve section was simulated by Equation 4 or Equation 5 (described in the Methods section). If the parameters of *N_0_*, *λ*, and *r* were taken as constants, Equation 4 was an exponential function; the value of *λN_0_*-*r* determined whether the curve was convexly upward, concavely downward or straight for this exponential function (Figure S2). In the case of *λN_0_*-*r*<0, the curve of this exponential function was convexly upward, the degree of convexly upward was greater with larger values of transcription rate coefficients if RNA degradation rate coefficients and *N_0_* were stable (Figure 3b1, Figure 3b2, and Figure 3b3), the degree of convexly upward was greater with smaller values of RNA degradation rate coefficients if transcription rate coefficient and *N_0_* were stable (Figure 3b6 and Figure 3b7), and the degree of convexly upward was greater with smaller values of *N_0_* if transcription and RNA degradation rate coefficients were stable (Figure 6f). In the case of *λN_0_*-*r*>0, the curve of this exponential function was concavely downward, the curve was less concavely downward with larger values of transcription rate coefficients if RNA degradation rate coefficients and *N_0_* were stable (Figure 3b2, Figure 3b3, and Figure 3b4), the curve was less concavely downward with smaller values of RNA degradation rate coefficients if transcription rate coefficient and *N_0_* were stable (Figure 3b5, Figure 3b6, and Figure 3b7), and the curve was less concavely downward with smaller values of *N_0_* if the transcription and RNA degradation rate coefficients were stable (Figure 6f). The combinations of different values of *λ* and *r*, and *N_0_* corresponded to the shape changes of local parts of the RNA level curves.

If the shapes of local parts of RNA level curves for direct simulation were smooth and monotonic, then these parts could be simulated well (Figure 6a, Figure 6b, Figure 6c, Figure 6d, and Figure 6e) because the curves of Equation 4 and Equation 5 were smooth and monotonic. There were four basic shapes of the curves of the simulation equations (Figure S2), and RNA level fluctuation data other than these four basic shapes could not be simulated well (Figure 3c and Figure 6f). A longer window length meant a higher possibility that a local part taken as a simulation unit of an RNA level curve was not smooth and not monotonic, leading to worse fitting (Figure 1a). The minimum window length was three, and the three points could form several shapes of the curve (Figure S3). Compared with the basic shapes of the curves of the simulation equations and the shapes of three-point curves (Figure S2 and Figure S3), three-point curves with monotonically increasing and convexly upward (Figure S3b), monotonically decreasing and concavely downward curves (Figure S3f), or horizontal lines (Figure S3d) could be simulated well; the three-point curves with monotonically increasing and concavely downward (Figure S3c) or monotonically decreasing and convexly upward (Figure S3e) seemed to be differently simulated; the three-point curves forming crests or valleys (Figure S3a) were most difficult to simulate by Equation 4 or Equation 5 (Figure 3c and Figure 6f). If more parts with monotonicity were added into the whole curves, the effects of worse fitting by crests and valleys could be diluted (Figure 6g, Figure 6h, Figure 6i, and Figure 6j). This could partly explain why the larger number of sampling time points generally resulted in better fitting indices (Figure 5). Simulated by the simulation curves with convexly upward or concavely downward shapes, which represented high transcription and degradation rate coefficients, the three-point curves with a crest, a valley, an increasing line followed by horizontal line, or a decreasing line followed by horizontal line have high transcription rate coefficients and high RNA degradation rate coefficients, meaning they have high estimation values of gross transcription rates and RNA degradation rates. The biological explanation might be that to reach appropriate RNA amounts in cells, an excessively high RNA amount caused by high transcription rates leads to high degradation rates and an excessively low RNA amount caused by high RNA degradation rates leads to high transcription rates because an excessively low or excessively high amount of RNA is harmful to cells (Carper, 1982; Cole et al., 1992). Generally, the crests with higher heights were well fitted by the simulation curves with higher transcription coefficients (Figure 3b1, Figure 3b2, and Figure 3b3), and the valleys with deeper depths were well fitted by the simulation curves with higher degradation coefficients (Figure 3b5, Figure 3b6, and Figure 3b7).

The shape of a local part of an RNA level curve was sometimes fitted by simulation results with different combinations of transcription rate coefficients and RNA degradation rate coefficients. For example, when *λN_0_*-*r*=0, the curve was a line:

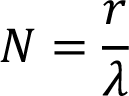

and different combinations of *r* and *λ* could achieve the same result. This produced a problem because *r* and *λ* determined the values of the gross transcription and RNA degradation rates, which meant that the same curve could be estimated for different gross transcription and RNA degradation rates, and the differences might be large. To solve this problem, the program of the present algorithm selected values of *r* and *λ* from small to large, and larger values could not replace smaller values if the simulation results were the same. Then, the smallest values were selected to estimate the related rates of the same curve. Whether this treatment conforms to biological laws is worth considering. Using a smaller metabolic RNA rate to obtain the same amount of RNA seems to be more resource efficient. Moreover, a slight change in the RNA level curve may cause a large change in the *λ* value in the simulation results, especially in a case with a concavely downward curve. Taking Figure 3b7 as an example, the first curve with a *λ* value of 0 and the fourth curve with a *λ* value of 0.012 (a total difference of 0.012) could be clearly separated, but under the same display multiple, the last curve with a *λ* value of 6.908 and the last fourth curve with a *λ* value of 4.084 (a total difference of 2.824) were difficult to distinguish. In the case of the exhaustive search method used in this study, reducing the value of the search step size might partially solve this problem; the search step size of *λ* is approximately equal to 0.007, which would be sufficient to obtain an acceptable *λ* value. A smaller search step size will obtain better results, but this requires a computer with higher computing power.

### 2.3 Effects of sampling time intervals

The design of sampling time intervals is important for time series gene expression experiments, and a time interval that is too large can easily produce erroneous experimental results (Xu and Asakawa, 2021). In the present study, the effects of sampling time intervals generally changed the shapes of RNA level curves and the estimation of gross transcription rates, depending on the specific RNA level curves.

In the RNA levels with high frequencies of crests and valleys, if the enlarged sampling time interval was larger than the width of the crest or the valley, the longer sampling time intervals could remove some details of fluctuations, resulting in smoother RNA level curves. This is equivalent to a reduction in the resolution of an image, resulting in ignoring the information for the image details. If the crests and valleys of the RNA level curves were skipped, longer sampling time intervals might lead to smaller gross transcription rates. For example, in Figure 4b, when the sampling time interval was extended from 10 to 20 minutes, some crests and valleys in the experimental data curve were missed, and the gross transcription rate dropped by 70%; in the zebrafish AF277172 gene, when the sampling time interval was extended from 1.3-18 hours to 21-24 hours, the gross transcription rate dropped by over 99% (Figure 4e); in the European eel Cathf gene, the gross transcription rate dropped by over 99% with an extended sampling time interval (Figure 4f). However, extending sampling time intervals did not always result in a decrease in the gross transcription rate of the RNA level fluctuation data. In the time length of three continuous sampling time points in the present study, the gross transcription rate was determined by the values of RNA abundance of the three time points. When the RNA abundances of the sample points in the treatment with the extended sampling time intervals were the same as those without extended sampling time intervals, the gross transcription rates could be the same, although the extended sampling time interval treatments ignored some crests and valleys (Figure S4a). If the sampling time points with higher RNA abundance were selected in the treatment with the extended sampling time intervals, longer sampling time intervals might result in higher gross transcription rates (Figure S4b). If the sampling time points with lower RNA abundance were selected in treatments with extended sampling time intervals, longer sampling time intervals might result in lower gross transcription rates (Figure S4c).

When the RNA level curve moved smoothly upward or downward without any crests or valleys, longer sampling time intervals generally made the RNA level curves less curved, resulting lower or higher estimated values of gross transcription rates, depending on the specific cases. If the RNA level curve had a convexly increasing monotonicity (Figure S5a), it was simulated by a convexly upward model curve (Figure S2a). Longer sampling time intervals made RNA level curves less curved and then made them be well fitted by less curved simulation curves, resulting in lower gross transcription rates of the sampling time interval from 10 minutes to 40 minutes but higher gross transcription rates of the sampling time interval from 40 minutes to 80 minutes. If the RNA level curve had a concavely increasing monotonicity (Figure S5b), it was simulated by an upward straight line in the model (Figure S2d). Longer sampling time intervals made RNA level curves less curved and changed the slope of the simulated lines; the gross transcription rates decreased with increasing sampling time intervals from 10 minutes to 40 minutes but increased with increasing time intervals from 40 minutes to 80 minutes. If the RNA level curve had convexly decreasing monotonicity (Figure S5c), it could only be simulated by the concavely downward model curve (Figure S2b), and longer sampling time intervals made the descending model curves more concave, with worse-fitting R-squared and MdAPE indices, resulting in lower gross transcription rates. If the RNA level curve had concavely decreasing monotonicity (Figure S5d), it was also simulated by the concavely downward model curve (Figure S2b); longer sampling time intervals generally resulted in lower gross transcription rates.

In a unimodal curve of RNA level, there was only one crest or valley, and longer sampling time intervals could make the local RNA level curve within a moving window more curved, thus resulting in higher gross transcription rates. In the case of the unimodal valley curve of the RNA level (Figure S6a), a valley with a depth of 100 units within the moving window resulting in the maximum transcription coefficients was formed by the sampling time points from the 70^th^ to the 90^th^ minute when the sampling time interval was 10 minutes, a valley with a depth of 1600 units within the moving window resulting in the maximum transcription coefficients was formed by the sampling time points from the 40^th^ to 120^th^ minute when the sampling time interval was 40 minutes, and a valley with a depth of 6400 units within the moving window resulting in the maximum transcription coefficients was formed by sampling time points from the 0^th^ to 160^th^ minute when the sampling time interval was 80 minutes; therefore, longer sampling time intervals led to higher gross transcription rates. The same effect was observed for longer sampling time intervals in the unimodal crest curve of the RNA level (Figure S6b). Similarly, in the analysis of yeast YBR067c (Figure 4d), when the sampling time interval was 10 minutes, a local valley with a depth of 104 units was formed by the sampling time points from the 120^th^ to the 140^th^ minute, and the transcription coefficients from the 100^th^ to the 140^th^ minute were 3.6∼45.8; however, when the sampling time interval extended to 20 minutes, a local valley with a depth of 155 units was formed by the sampling time points from the 100^th^ to the 140^th^ minute, and the transcription coefficient from the 100^th^ to the 140^th^ minute was 1182.0; thus, the change in transcription coefficients in this local valley greatly contributed to the change in the global gross transcription rates.

If the origin RNA level curve was a straight line, longer sampling time intervals could not change the shape of the line because two points already define a straight line, but the effects on the gross transcription rates depended on the direction of the line. In the case of the upward straight line of the RNA level, the data could be simulated by the model curve of the oblique upward straight line (Figure S2d), and the longer sampling time intervals could not change the estimation of the gross transcription rate. If the RNA level curve is a horizontal line, the data could be simulated by the model curve of the horizontal line (Figure S2c), and longer sampling time intervals could not change the estimation of gross transcription rates. In the case of the downward straight line of the RNA level, longer sampling time intervals could not change the shape of the RNA level line, but the data were simulated by the concavely downward curve of the model (Figure S2b), resulting in lower estimated gross transcription rates.

Thus, the results of the present study showed that the longer the sampling time interval was, the higher the possibility of affecting the shapes of experimental RNA level curves and the estimations of gross transcription rates. Which range of sampling time interval could be used to obtain a more accurate measurement of RNA level fluctuation curves and estimated gross transcription rates? According to fractal theory, within a certain range, the smaller the sampling time interval is, the more accurate the RNA level curve that can be obtained, with more accurate estimated values of gross transcription rates. Smaller sampling time intervals, however, mean more work, higher cost, and more time commitment, which is unacceptable for some experimental designs. Theoretically, if every RNA molecule transcribed during the experiment could be detected, the RNA level curve would likely contain a more complete amount of transcriptional information. That is, the sampling time intervals were shorter than the lifespans of RNAs. If half-lives of RNA can be considered representative lifespans, the lifespans of mRNAs in prokaryotic cells range from approximately 1∼50 minutes (Cai and Winkler, 1993; Steiner et al., 2019), and the lifespans of mRNAs in eukaryotic cells range from 10 minutes to 48 hours (Petersen et al., 1976; Deng et al., 2013; McManus et al., 2015; Yamada and Akimitsu, 2019); this information may help to guide the arrangement of sampling time intervals of related experiments. The lifespan of RNAs might be renewed by developing detection technology (Baudrimont et al., 2017), which affects the design of the sampling time interval. When the RNA level fluctuation data are obtained, the effects of longer sampling time intervals can be analyzed, as in this study, but the effects of sampling time intervals shorter than those in actual experiments are difficult to predict. Research on this topic deserves further development.

## 3 Conclusion

RNA level fluctuation data, in which sampling time intervals influence the resolution of the dynamic trends of continuous gene expression, are accumulating with the development of RNA quantitative assay technology, and estimating transcription rates from these data is an urgent task for downstream transcriptomic research. The present study can directly estimate gross transcription rates from RNA level fluctuation data. Among the 384 yeast genes in this study, the 20 genes with the highest gross transcription rates played main roles in cell division regulation, DNA replication, biosynthesis of proteins and fatty acids, and material transport, which were slightly different from the results of the analysis with metrics such as raw gene expression data and gene expression averages. The results for analyzing gross transcription rate × gene length showed that the gene consuming the most gene expression resources functioned in fatty acid biosynthesis and protein transport. For some studies, the results obtained by analyzing the data in terms of gross transcription rate may provide important information. The acquisition of gross transcription rates in routine gene expression experiments will provide more chances to engage further transcriptomic research.

Special shapes of RNA level curves were linked with large changes in local gross transcription rates. The transcription and RNA degradation rates were both higher in the local parts with narrow crests or valleys of RNA level curves. Different sampling time intervals could affect the shapes of RNA level curves and the affect estimations of gross transcription rates. Therefore, how to use the available information to design the sampling time intervals in time course gene expression experiments needs to be highly regarded. Although these results represent the analyses performed with the present algorithm, sampling time intervals may generally affect gene expression phenomena.

The present algorithm was developed for analyzing RNA level fluctuation data, and it has the potential to be used in other related data, such as data for other biomolecules in cells and RNAs in different environments. Further related research will be worthwhile.

## 4 Methods

### 4.1 Description of the model

#### a. The basic unit of the model

Transcription and RNA degradation have different functional relationships with RNA abundance (Gorini and Maas, 1957; Zeisel et al., 2011; Palumbo et al., 2015; Yamada and Akimitsu, 2019). Transcription was as follows:

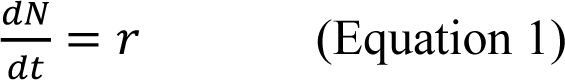

where *N* is RNA abundance, *t* is time, and *r* is the transcription rate coefficient. RNA degradation was as follows:

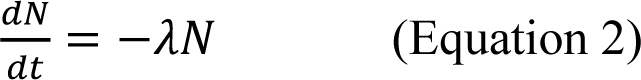

where *λ* is the degradation rate coefficient. RNA abundance equaled transcription minus degradation, described as follows:

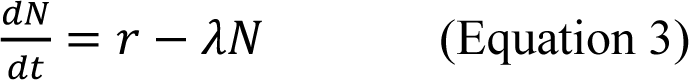

By solving differential Equation 3, the dynamic change in RNA abundance over time was as follows:

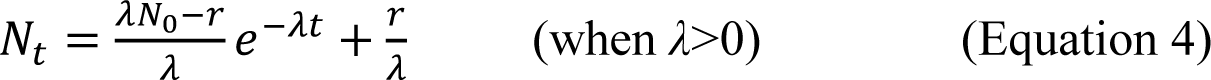

where *N_t_* is the RNA abundance at time t, and *N_0_* is the RNA abundance at time 0. Equation 4 describes the RNA abundance dynamics only under the situation that the degradation rate is larger than zero, but the situation in which the degradation rate is zero can exist. If the degradation rate is zero, the dynamic change in RNA abundance over time was obtained by solving differential Equation 1:

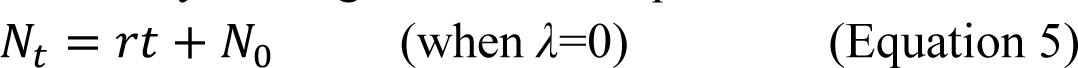

#### b. The moving window

In Equation 4 or Equation 5, the transcription and degradation rate coefficients were constants, but during the experiment time periods, transcription and degradation rate coefficients usually changed, leading to RNA level fluctuations; thus, it was difficult to directly use Equation 4 or Equation 5 to simulate RNA levels over the experiment time. To solve this problem, we used a moving window to set several continuous sampling time points as a simulation unit, and Equation 4 or Equation 5 was used to simulate the data within the moving window. When the simulation analysis of a previous moving window was completed, the moving window moved a sampling time point to analyze the next moving window (Figure S7).

#### c. Assignment of initial values

During the simulation of data in each moving window using Equation 4 or Equation 5, the values of r and λ were set with a certain step size, and the exhaustive search method was used to select the *r* and *λ* values that made the simulated curve fit the experimental value curve the best. In each moving window, the sum of squared differences between the predicted values and the experimental values at each sampling time point was used as an indicator of the degree of fit. The maximum value, the minimum value and the step size of *r* and *λ* could be set according to the specific experimental data analyzed. In the present study, the minimum value of *r* was zero, the maximum value of *r* was double the maximum RNA abundance in the experimental data, and the step size of *r* was set as:

Step size of *r* = (Maximum value of *r* - Minimum value of *r*)/10000.

The doubling of the maximum RNA abundance resulted from an assumption that the RNA amount transcribed at the sampling time point just before the sampling time point of the maximum RNA abundance at most degraded 50% to the next sampling time point to be the maximum RNA abundance immediately. To set the maximum value and minimum value of *λ*, differential Equation 2 was solved as:

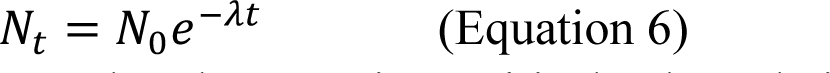

where *t* is time, *N_t_* is the RNA abundance at time t, *λ* is the degradation rate coefficient, and *N_0_* is the RNA abundance at time 0. The minimum value of *λ* was set to mean that the RNA amount transcribed at a certain sampling time point was maintained at 100% of the value at the next sampling time point, and it was calculated as follows:

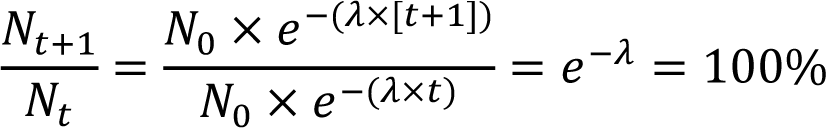

Thus, the minimum value of *λ* was zero. The maximum value of *λ* was set to mean that the RNA amount transcribed at a certain sampling time point was maintained at only 0.1% of the next sampling time point, and it was calculated as follows:

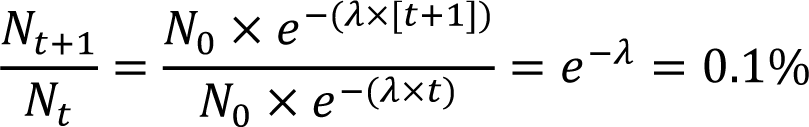

Thus, the maximum value of *λ* was 6.90776. The step size of *λ* was set as:

The step size of *λ* = (The maximum value of *λ* - The minimum value of *λ*)/1000.

#### d. Modeling in the moving windows

In a moving window, the predicted RNA abundance *N_t_* could be calculated by Equation 4 or Equation 5 at each sampling time point. If a sampling time point was covered by more than one moving window, the mean predicted RNA abundance at this sampling time point was calculated as follows:

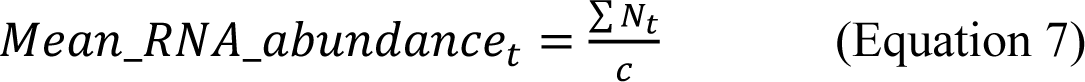

where *t* is time, *Mean_RNA_abundance_t_* is the mean predicted RNA abundance at time *t*, *N_t_* is the predicted RNA abundance at time *t* in a moving window, and *c* is the number of the moving window in which the sampling time point at time *t* is covered. When the mean predicted RNA abundance at all sampling time points was obtained, the fitting index between the predicted RNA abundance and experimental data was calculated. The R-squared and MdAPE values were herein set as the fitting indices.

#### e. Calculating gross transcription rates

The net RNA amount without degradation could be directly calculated as follows if *r* was obtained by the simulation in a moving window:

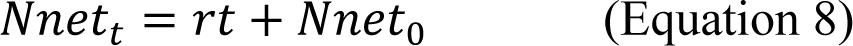

where *t* is time, *Nnet_t_* is the amount of RNA without degradation at time *t*, *r* is the transcription rate coefficient, and *Nnet*_0_ is the amount of RNA without degradation at time 0. The transcription amount at time *t* in a moving window was calculated as follows:

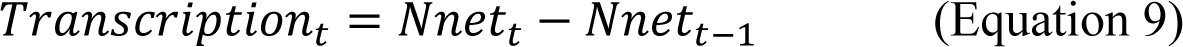

where *Transcription_t_* is the transcription amount at time *t*, *Nnet_t_* is the RNA amount without degradation at time t, and *Nnet_t-1_* is the RNA amount without degradation at time t-1. If a sampling time point was covered by more than one moving window, the mean transcription amount at time *t* was calculated as follows:

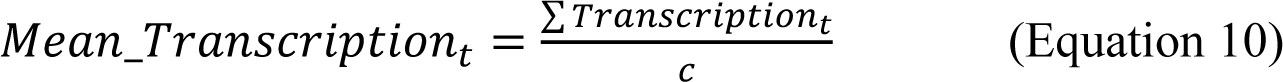

where *Mean_Transcription_t_* is the mean transcription amount at time *t*, and *c* is the number of the moving window in which the sampling time point at time *t* was covered. The cumulative transcription was calculated as follows:

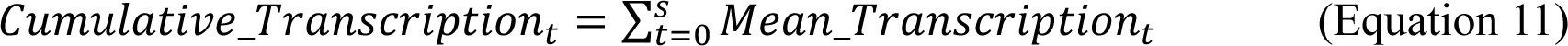

where *Cumulative_Transcription_t_* is the cumulative transcription amount during time *t*, *s* is a specified point of time, and *Mean_Transcription_t_* is the mean transcription amount at time *t*. The gross transcription rate was calculated as follows:

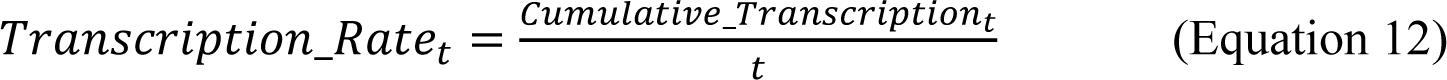

where *Transcription_Rate_t_* is the transcription rate during time *t*, and *Cumulative_Transcription_t_* is the cumulative transcription amount during time *t*.

#### f. Calculation of the RNA degradation rates

The RNA degradation amount could not be calculated by Equation 6 because RNA degradation depended on the RNA concentration and *N_0_*, which changed due to transcription. The RNA degradation amount in a moving window was calculated as follows:

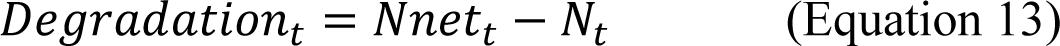

where *t* is time, *Degradation_t_* is the RNA degradation amount at time *t*, *Nnet_t_* is the RNA amount without degradation at time t, and *N_t_* is the RNA abundance at time t. If a sampling time point was covered by more than one moving window, the mean RNA degradation amount at time *t* was calculated as follows:

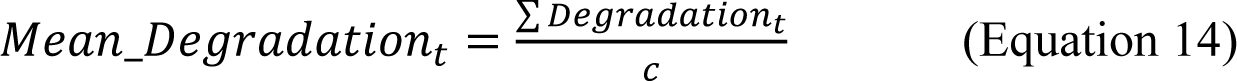

where *Mean_Degradation_t_* is the mean RNA degradation amount at time *t*, *Degradation_t_* is the RNA degradation amount at time *t* in a moving window, and *c* is the number of the moving window in which the sampling time point at time *t* was covered. The cumulative RNA degradation was calculated as follows:

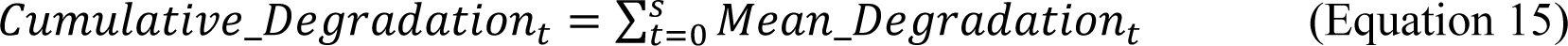

where *Cumulative_Degradation_t_* is the cumulative RNA degradation amount during time *t*, *s* is a specified time point, and *Mean_Degradation_t_* is the mean RNA degradation amount from time *t*. The RNA degradation rate was calculated as follows:

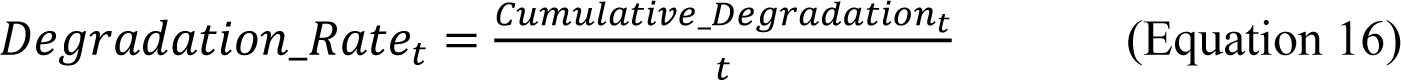

where *Degradation_Rate_t_* is the RNA degradation rate during time *t*, and *Cumulative_Degradation_t_* is the cumulative RNA degradation amount during time *t*.

### 4.2 Dataset

The data of time-series gene expression experiments of the yeast *Saccharomyces cerevisiae* (Cho et al., 1998; Yeung et al., 2001), zebrafish *Danio rerio* (Mathavan et al., 2005), and European eel *Anguilla* (Bolliet et al., 2017) were used as the study objects (Table S2). The experimental data of yeast were measured by the oligonucleotide array method, the experimental data of zebrafish were measured by the microarray method, and the experimental data of European eel were measured by RT‒PCR.

### 4.3 Calculation of gross transcription and RNA degradation rates in the datasets

The experimental RNA levels of the *S. cerevisiae* YML021C gene were used to determine the appropriate length of the moving window, and the fitting indices of R-squared and MdAPE values were used as criteria for judging which window length was the best. To illustrate the generality of the result of the best moving window length, the data of the genes of the first three maxima of mean RNA abundance, the three medians of mean RNA abundance, and the first three minima of mean RNA abundance among the 384 *S. cerevisiae* gene dataset were selected to determine the suitable length of the moving window.

The gross transcription and RNA degradation rates of the genes in the yeast, zebrafish, and European eel datasets were calculated by using this algorithm. Clustering of gross transcription rates, consumption rates of nucleotides (e.g., gross transcription rate × gene length), mean RNA abundance, and raw RNA abundance data with the yeast dataset of 384 genes was analyzed. Clustering of the one-time-point RNA abundance of 384 yeast genes was also analyzed.

The experimental RNA abundance data of the yeast YER111c and YELo66W genes were used to analyze the effects of the shapes of RNA level curves on the calculation of gross transcription rate. The datasets of different transcription coefficients and different RNA degradation coefficients under the simulation background were analyzed to help explain the effects of the shapes of RNA level curves.

The experimental RNA level fluctuation data of the yeast YLR079w gene, yeast YKL066W gene, yeast YDL198c gene, yeast YBR067c gene, zebrafish AF277172 gene, and European eel Cathf gene were used to analyze the effects of sampling time intervals on the calculation of gross transcription rates. The data of longer sampling time intervals were set by selecting the raw sampling time point data for every time interval designed.

The experimental RNA level fluctuation of the yeast YJL194w gene was used to study the effects of the number of sampling time points on the calculation of gross transcription rates and RNA degradation rates, and the fitting indices of the R-squared and MdAPE values were used as criteria for comparison of the results of different numbers of sampling time points. To illustrate the generality of the results of different numbers of sampling time points, the data of the genes of the first three maximum mean RNA abundances, the three median mean RNA abundances, and the first three minimum mean RNA abundances among the 384 yeast gene datasets were selected to study the effects of the number of sampling time points.

The datasets of linear function, exponential function, fractional function, parabolic function, sine function, and periodic fluctuation curves under the simulation background were used to analyze the ability of this algorithm to simulate curves of different shapes. The dataset with different values of *N_0_* in Equation 4 under the simulation background was analyzed to help explain why the fluctuations with short circling lengths were difficult to simulate.

### 4.4 Programming

The simulation processes were translated into the C++ language (provided upon request). The R language was used to draw the curves of the simulation outputs (provided upon request).

## Acknowledgments

This work was supported by JST SPRING, Grant Number JPMJSP2108.

## Figures

## Supplemental tables

**Table S2.**
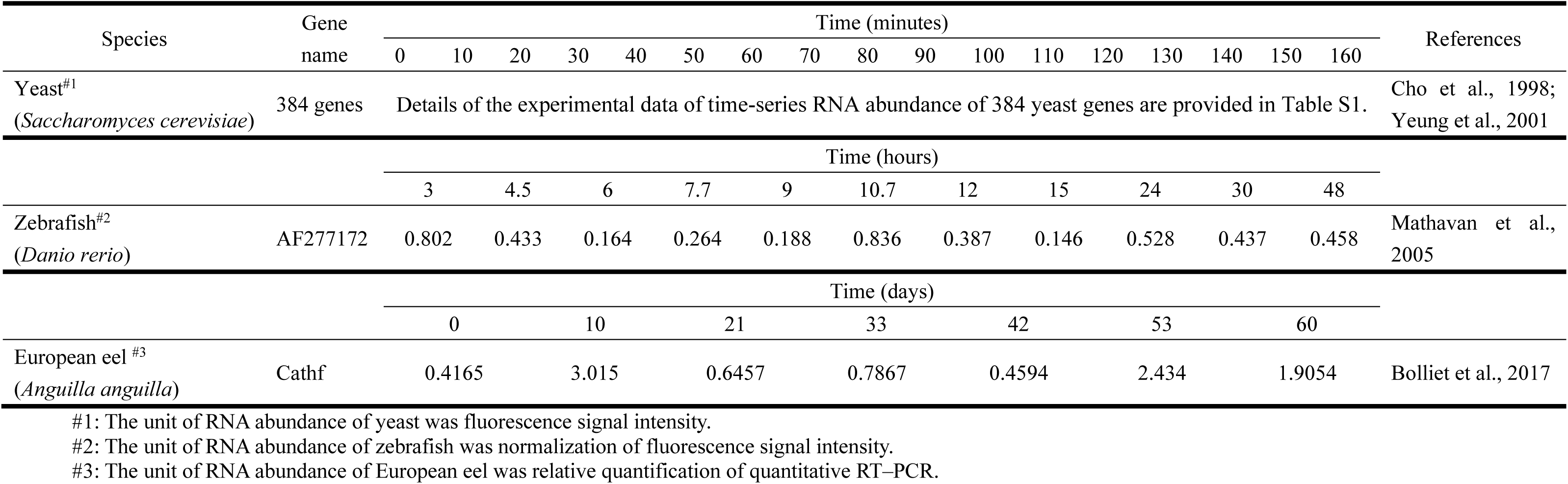
Experimental dataset of RNA level fluctuation used in the present study

## Supplemental figures

**Figure S1.**
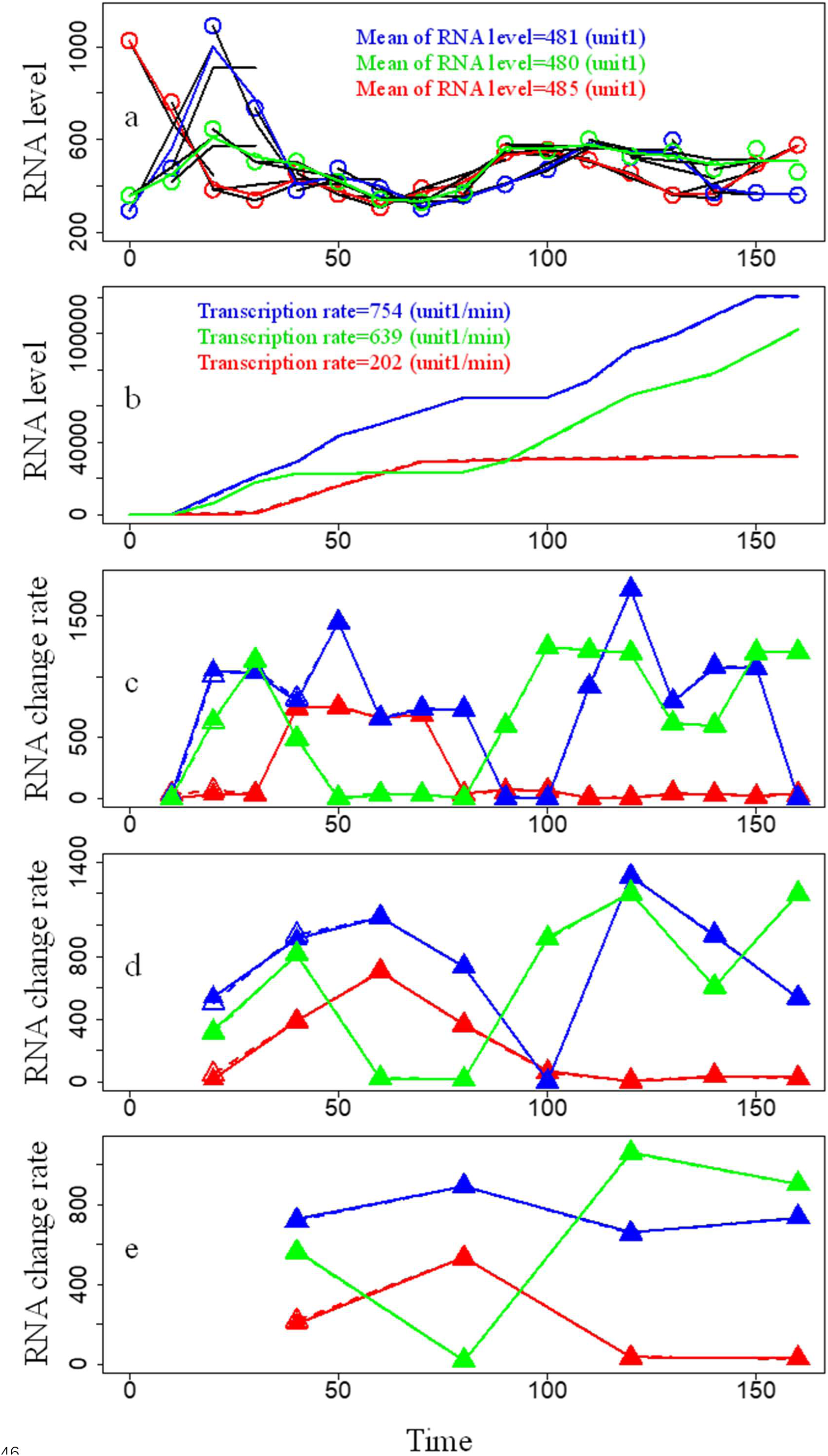
Distinguishing gene expression intensities of three genes with RNA level fluctuations by the present algorithm. The experimental data of yeast RNA levels were from the literature (Cho et al., 1998; Yueng et al., 2001). The results of the yeast YDL124w gene are shown in red, the results of the yeast YKL045w gene are shown in blue, and the results of the yeast YGL200c gene are shown in green. Transcription rates denoted gross transcription rates. **a.** RNA level fluctuations of the three yeast genes with similar mean RNA levels. The circles represent experimental data points. The black curves represent simulation curves of a moving window with a window length of three. Unit 1 is the fluorescence signal intensity. The x-axis is time, and the unit of time is minutes. The y-axis is the RNA level, and the unit is the fluorescence signal intensity. **b.** Accumulation of transcription and RNA degradation of the three yeast genes. The solid curves represent transcription accumulation. The dashed curves represent RNA degradation accumulation. Unit 1 is the fluorescence signal intensity. The x-axis is time, and the unit of time is minutes. The y-axis is the RNA level, and the unit is the fluorescence signal intensity. **c.** Gross transcription rates and RNA degradation rates calculated every ten minutes. The solid triangles represent data points of the gross transcription rate calculated every ten minutes. The hollow triangles represent data points of the RNA degradation rate calculated every ten minutes. Some gross transcription and RNA degradation rate points overlapped. The x-axis is time, and the unit of time is minutes. The y-axis is the gross transcription rate or RNA degradation rate, and the unit is the fluorescence signal intensity per minute (unit 1/min). **d.** Gross transcription rates and RNA degradation rates calculated every twenty minutes. The solid triangles represent data points of the gross transcription rate calculated every twenty minutes. The hollow triangles represent data points of the RNA degradation rate calculated every twenty minutes. Some gross transcription and RNA degradation rate points overlapped. The x-axis is time, and the unit of time is minutes. The y-axis is the gross transcription rate or RNA degradation rate, and the unit is the fluorescence signal intensity per minute (unit 1/min). **e.** Gross transcription rates and RNA degradation rates calculated every forty minutes. The solid triangles represent data points of the gross transcription rate calculated every forty minutes. The hollow triangles represent data points of the RNA degradation rate calculated every forty minutes. Some gross transcription and RNA degradation rate points overlapped. The x-axis is time, and the unit of time is minutes. The y-axis is the gross transcription rate or RNA degradation rate, and the unit is the fluorescence signal intensity per minute (unit 1/min).

**Figure S2.**
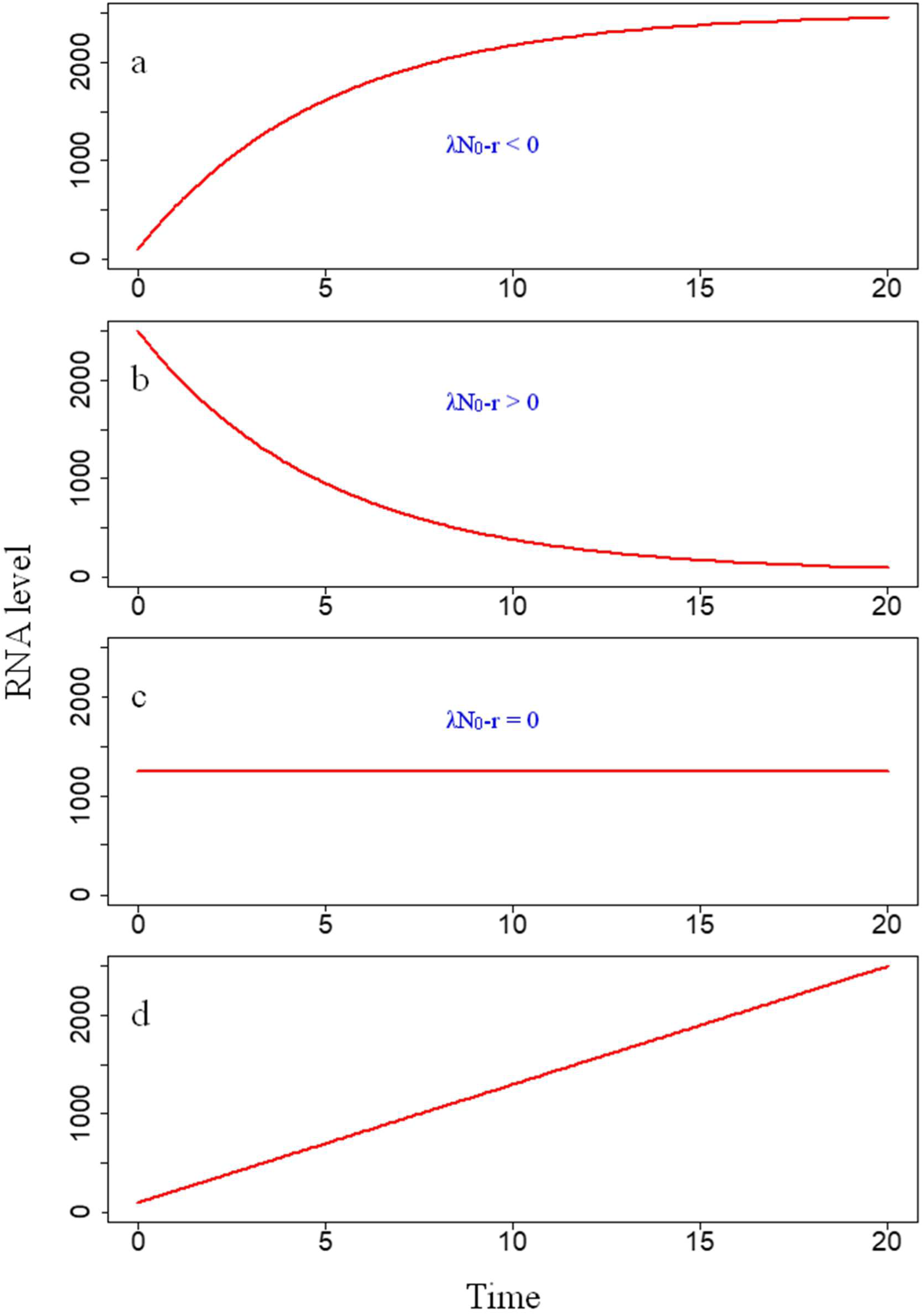
Typical shapes of the curves of basic equations in the algorithm of the present study. **a.** Convexly upward curve: 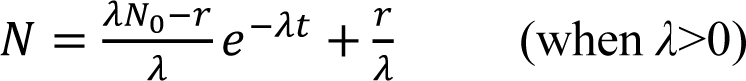. **b.** Concavely downward curve: 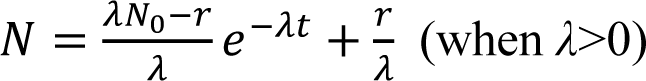. **c.** Horizontal line curve: 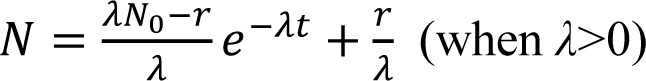. **d.** Oblique upward straight line curve: 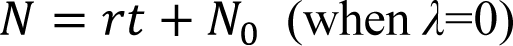.

**Figure S3.**
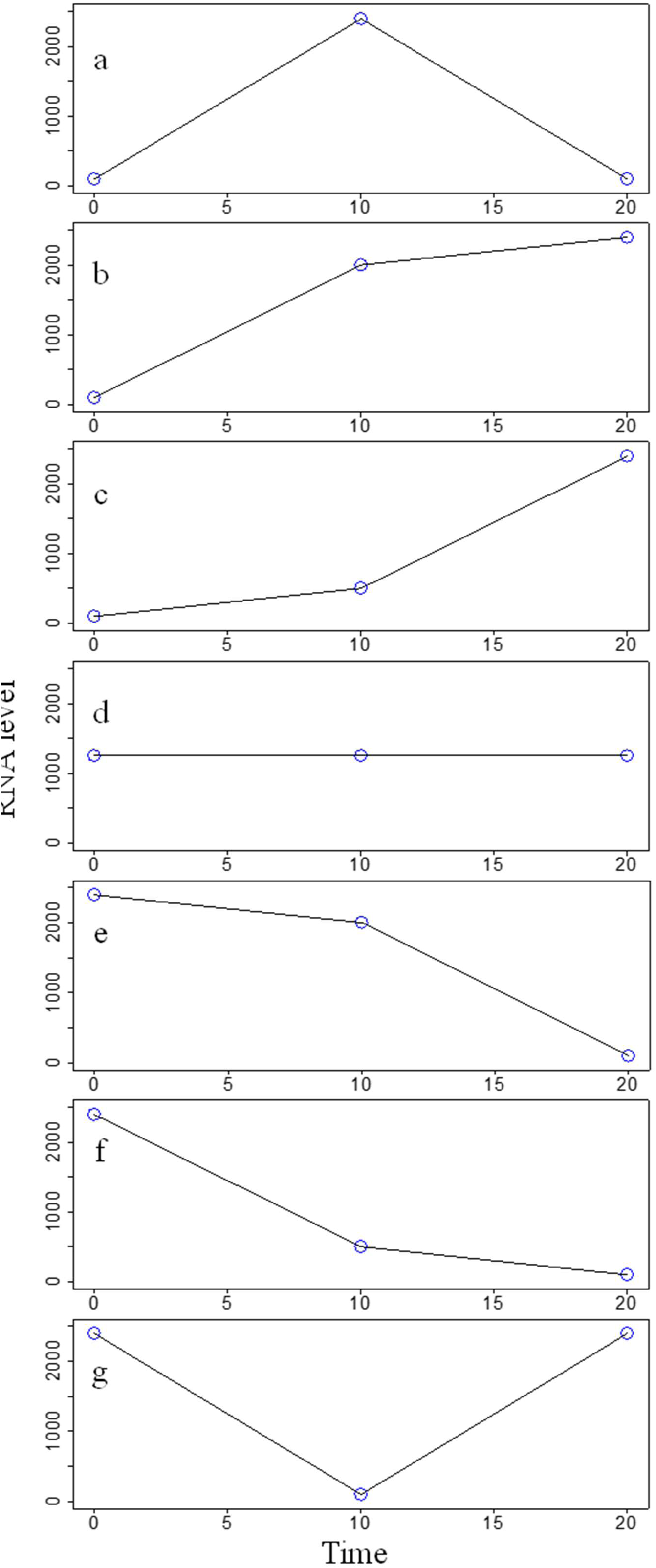
Typical shapes of the curves formed by three points. **a.** Crest. A curve with this shape could be simulated by the model curve with the shape in Figure S2a. **b.** Monotonically increasing and convexly upward curve. This type of curve includes an oblique upward straight line. A curve with this shape could be simulated by the model curve with the shape in Figure S2a and Figure S2d. **c.** Monotonically increasing and concavely downward curve. A curve with this shape could be simulated by the model curve with the shape in Figure S2a and Figure S2d. **d.** Horizontal line. A curve with this shape could be simulated by the model curve with the shape in Figure S2c. **e.** Monotonically decreasing and convexly upward curve. This type of curve includes an oblique downward straight line. A curve with this shape could be simulated by the model curve with the shape in Figure S2b. **f.** Monotonically decreasing and concavely downward curve. A curve with this shape could be simulated by the model curve with the shape in Figure S2b. **g.** Valley. A curve with this shape could be simulated by the model curve with the shape in Figure S2b.

**Figure S4.**
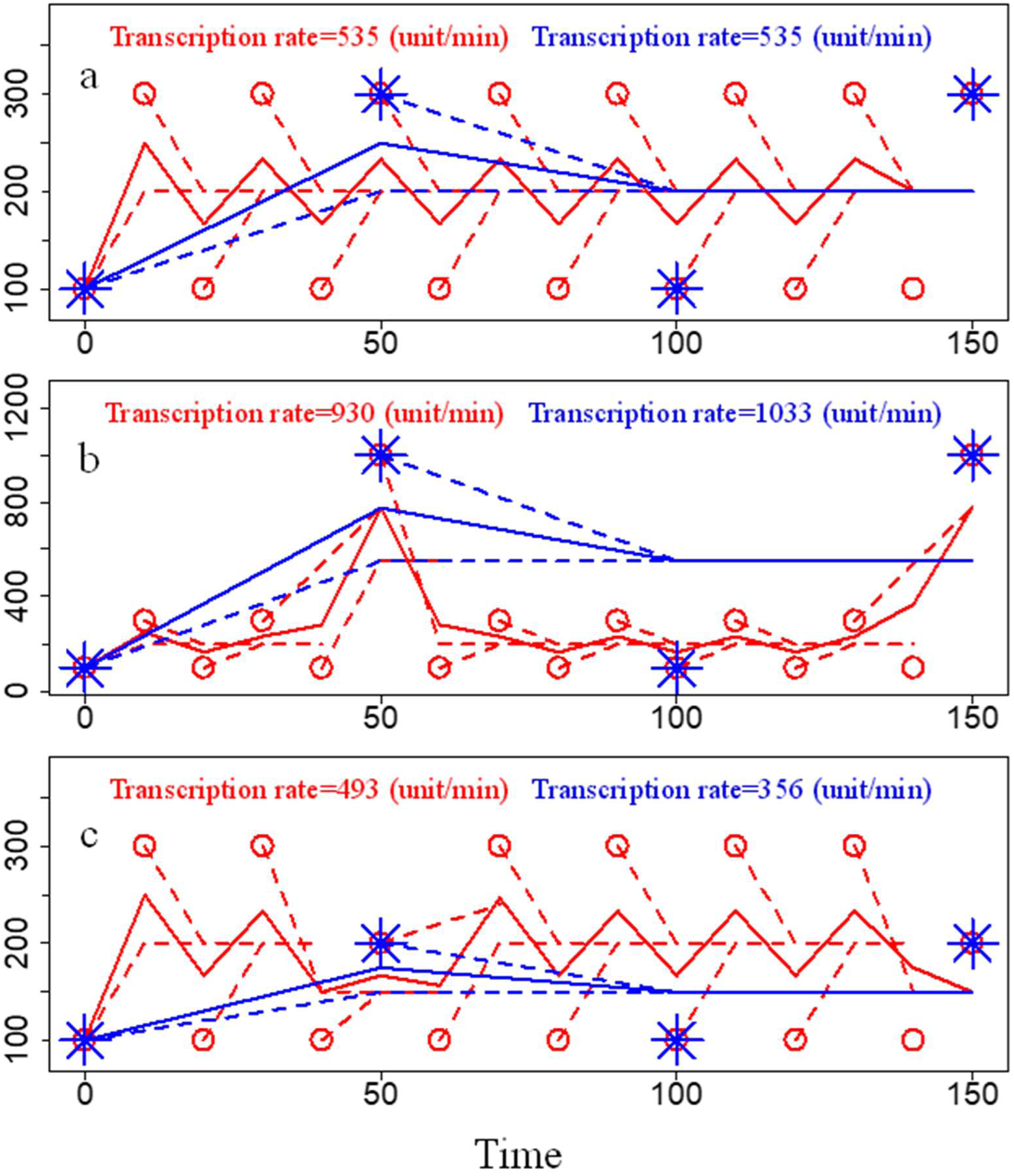
Different effects of expanding sampling time intervals on the estimations of gross transcription rates when the sampling time intervals are longer than circling length of RNA level fluctuations. The results with a sampling time interval of 10 minutes are shown in red, and the results with a sampling time interval of 50 minutes are shown in blue. The circles and asterisks are experimental data points. The dashed curves represent simulation curves of a moving window with a window length of three. The solid curves represent the mean RNA abundances of the points calculated in different moving windows. The x-axis is time, and the unit of time is minutes. The y-axis is the RNA level, and the unit is the fluorescence signal intensity. Transcription rates denoted gross transcription rates. **A.** No difference in gross transcription rates with longer sampling time intervals. In the results with a sampling time interval of 10 minutes, the RNA degradation rate was 534 units/min, the R-squared value was 0.526, and the MdAPE value was 0.278. In the results with a sampling time interval of 50 minutes, the RNA degradation rate was 534 units/min, the R-squared value was 0.438, and the MdAPE value was 0.250. **b.** Longer sampling time intervals resulted in higher gross transcription rates. In the results with a sampling time interval of 10 minutes, the RNA degradation rate was 926 unit/min, the R-squared value was 0.784, and the MdAPE value was 0.223. In the results with a sampling time interval of 50 minutes, the RNA degradation rate was 1032 units/min, the R-squared value was 0.438, and the MdAPE value was 0.338. **c.** Longer sampling time intervals resulted in lower gross transcription rates. In the results with a sampling time interval of 10 minutes, the RNA degradation rate was 493 units/min, the R-squared value was 0.594, and the MdAPE value was 0.236. In the results with a sampling time interval of 50 minutes, the RNA degradation rate was 356 unit/min, the R-squared value was 0.438, and the MdAPE value was 0.188.

**Figure S5.**
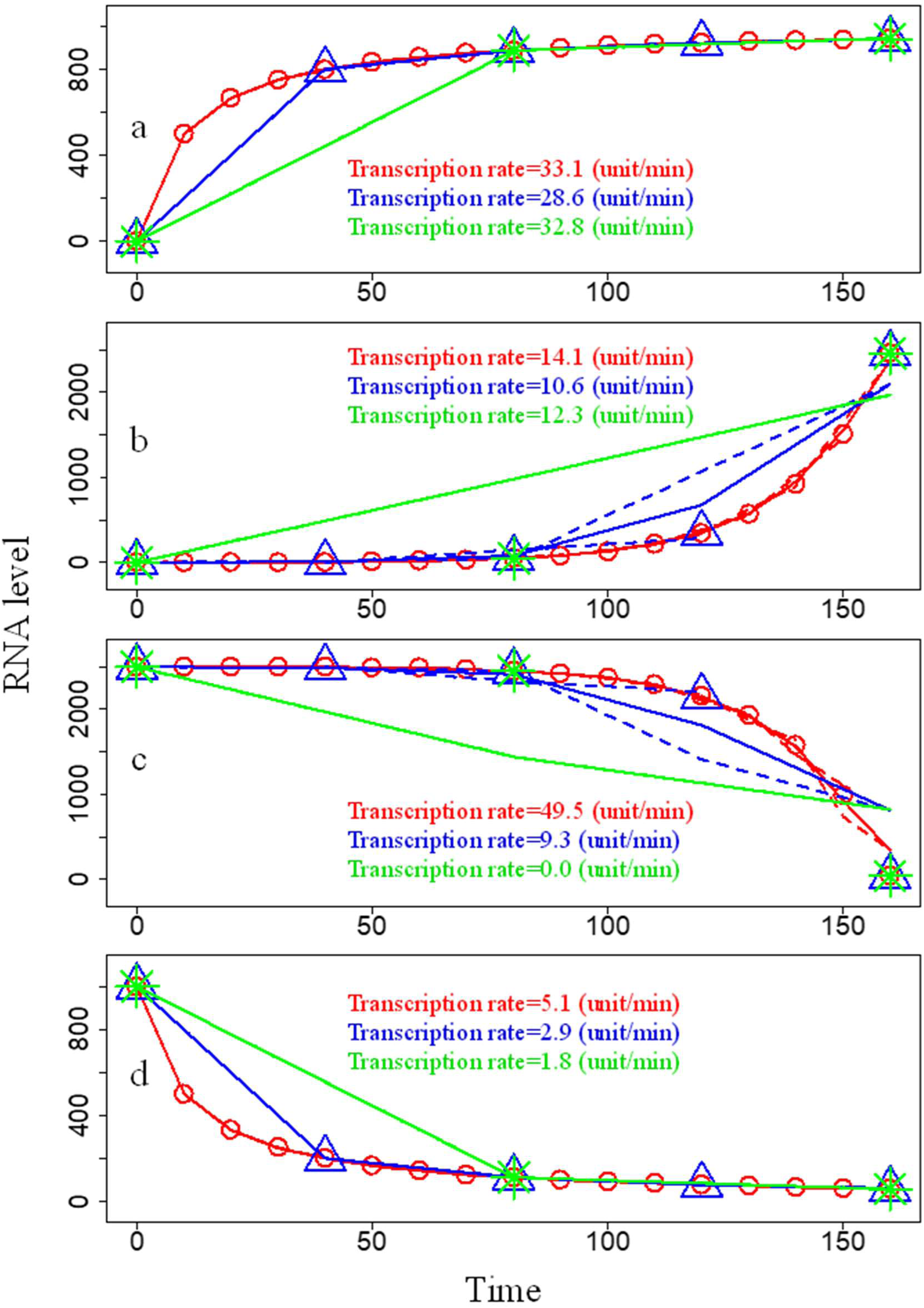
Effects of expanding sampling time intervals on the estimations of gross transcription rates in monotonic curves of RNA levels. The results with a sampling time interval of 10 minutes are shown in red, the results with a sampling time interval of 40 minutes are shown in blue, and the results with a sampling time interval of 80 minutes are shown in green. The circles, triangles, and asterisks are experimental data points. The dashed curves represent simulation curves of a moving window with a window length of three. The solid curves represent the mean RNA abundances of the points calculated in different moving windows. The x-axis is time, and the unit of time is minutes. The y-axis is the RNA level, and the unit is the fluorescence signal intensity. Transcription rates denoted gross transcription rates. **a.** Convexly increasing monotonic curve. In the results with a sampling time interval of 10 minutes, the RNA degradation rate was 27.2 unit/min, the R-squared value was 1.000, and the MdAPE value was 0.000. In the results with a sampling time interval of 40 minutes, the RNA degradation rate was 22.7 unit/min, the R-squared value was 1.000, and the MdAPE value was 0.000. In the results with a sampling time interval of 80 minutes, the RNA degradation rate was 26.8 units/min, the R-squared value was 1.000, and the MdAPE value was 0.000. **b.** Concavely increasing monotonic curve. In the results with a sampling time interval of 10 minutes, the RNA degradation rate was 0.1 unit/min, the R-squared value was 0.999, and the MdAPE value was 0.023. In the results with a sampling time interval of 40 minutes, the RNA degradation rate was 0.0 unit/min, the R-squared value was 0.947, and the MdAPE value was 0.577. In the results with a sampling time interval of 80 minutes, the RNA degradation rate was 0.0 unit/min, the R-squared value was 0.718, and the MdAPE value was 0.200. **c.** Convexly descending monotonic curve. In the results with a sampling time interval of 10 minutes, the RNA degradation rate was 61.5 unit/min, the R-squared value was 0.987, and the MdAPE value was 0.001. In the results with a sampling time interval of 40 minutes, the RNA degradation rate was 17.4 unit/min, the R-squared value was 0.843, and the MdAPE value was 0.017. In the results with a sampling time interval of 80 minutes, the RNA degradation rate was 10.4 unit/min, the R-squared value was 0.584, and the MdAPE value was 0.413. **d.** Concavely decreasing monotonic curve. In the results with a sampling time interval of 10 minutes, the RNA degradation rate was 11.0 unit/min, the R-squared value was 1.000, and the MdAPE value was 0.001. In the results with a sampling time interval of 40 minutes, the RNA degradation rate was 8.8 units/min, the R-squared value was 1.000, and the MdAPE value was 0.001. In the results with a sampling time interval of 80 minutes, the RNA degradation rate was 7.7 unit/min, the R-squared value was 1.000, and the MdAPE value was 0.007.

**Figure S6.**
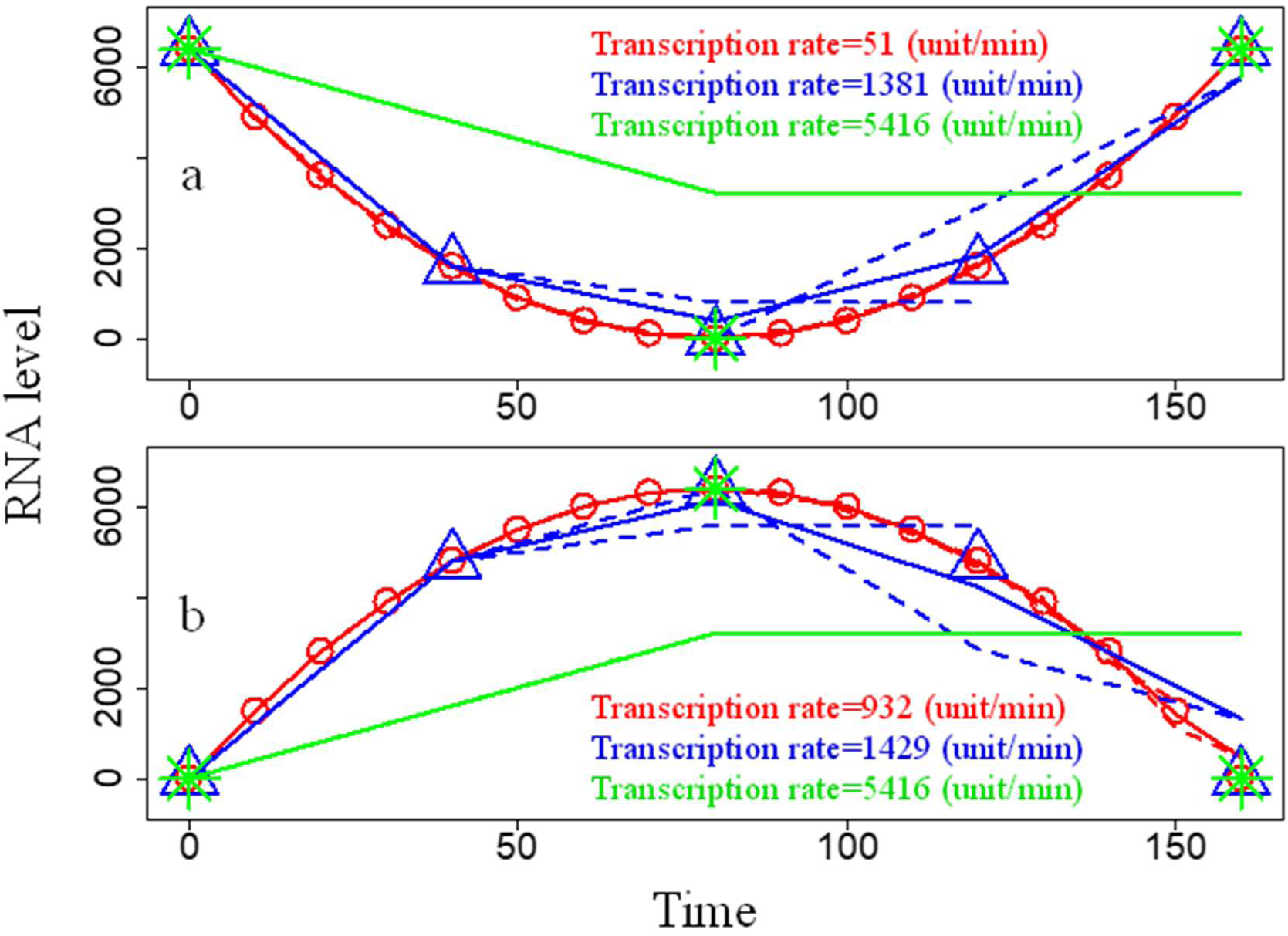
Effects of expanding sampling time intervals on the estimations of gross transcription rates in unimodal curves. The results with a sampling time interval of 10 minutes are shown in red, the results with a sampling time interval of 40 minutes are shown in blue, and the results with a sampling time interval of 80 minutes are shown in green. The circles, triangles, and asterisks are experimental data points. The dashed curves represent simulation curves of a moving window with a window length of three. The solid curves represent the mean RNA abundances of the points calculated in different moving windows. The x-axis is time, and the unit of time is minutes. The y-axis is the RNA level, and the unit is the fluorescence signal intensity. Transcription rates denote gross transcription rates. **a.** Parabolic curve with an upward opening. In the results with a sampling time interval of 10 minutes, the RNA degradation rate was 52 unit/min, the R-squared value was 1.000, and the MdAPE value was 0.006. In the results with a sampling time interval of 40 minutes, the RNA degradation rate was 1390 units/min, the R-squared value was 0.982, and the MdAPE value was 0.104. In the results with a sampling time interval of 80 minutes, the RNA degradation rate was 5436 units/min, the R-squared value was 0.250, and the MdAPE value was 0.500. **b.** Parabolic curve with a downward opening. In the results with a sampling time interval of 10 minutes, the RNA degradation rate was 927 unit/min, the R-squared value was 0.997, and the MdAPE value was 0.002. In the results with a sampling time interval of 40 minutes, the RNA degradation rate was 1412 units/min, the R-squared value was 0.940, and the MdAPE value was 0.042. In the results with a sampling time interval of 80 minutes, the RNA degradation rate was 5396 units/min, the R-squared value was 0.250, and the MdAPE value was 0.500.

**Figure S7.**
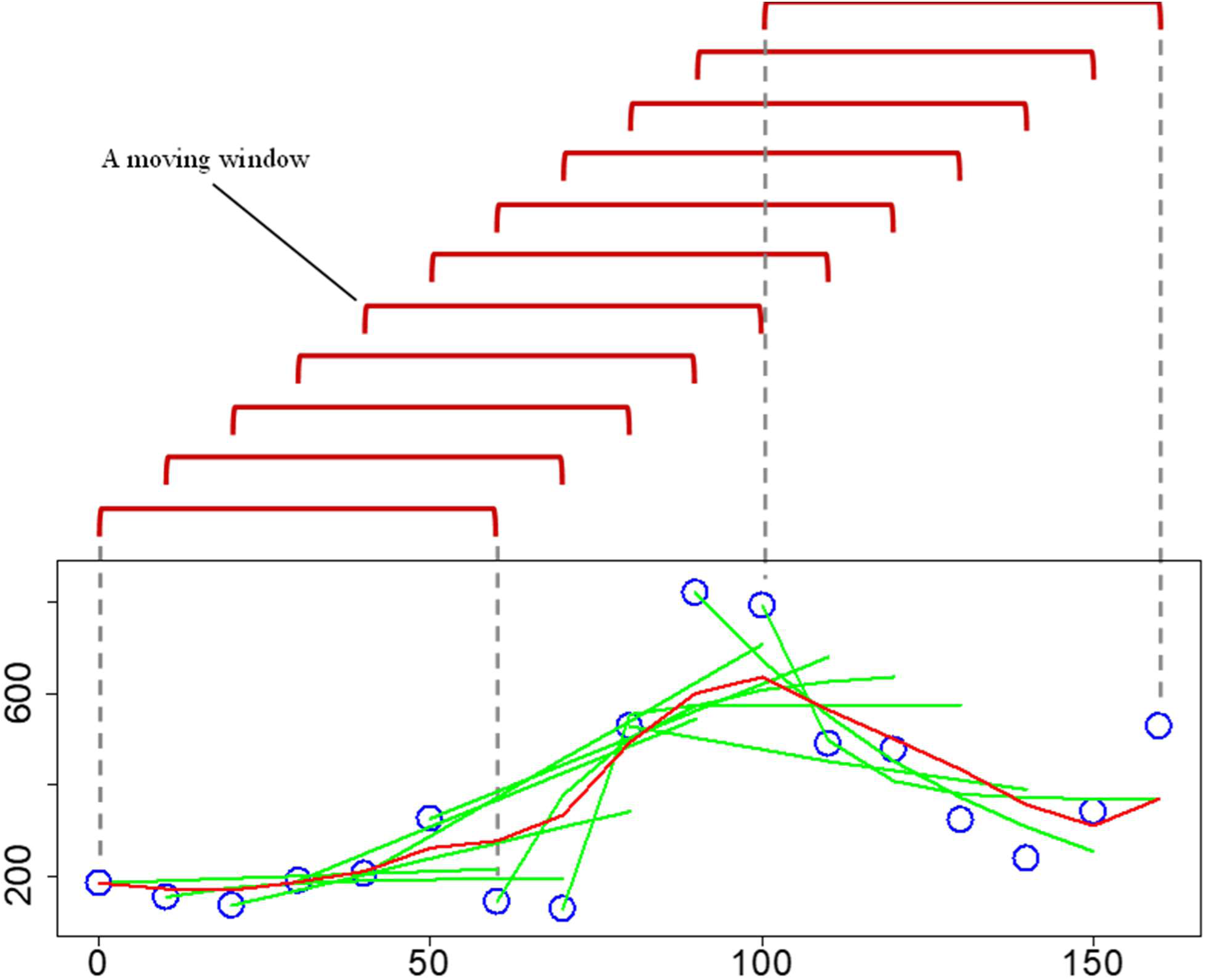
Moving window of the present algorithm. Blue circles are experimental data points. The brown half boxes indicate that there was a moving window (here, a window length of 7 was used as an example). The green curves represent simulation curves for 7 consecutive values in a moving window by using Equation 4 or Equation 5. The red curve represents the mean of the predicted values of the points within different moving windows. The x-axis shows the time, and the y-axis shows the RNA level.

## References

1. Alkallas R, Fish L, Goodarzi H, Najafabadi HS. Inference of RNA decay rate from transcriptional profiling highlights the regulatory programs of Alzheimer’s disease. Nat Commun. 2017 Oct 13;8(1):909. doi: 10.1038/s41467-017-00867-z. Erratum in: Nat Commun. 2018 Oct 31;9(1):4625.

2. Armingol E, Officer A, Harismendy O, Lewis NE. Deciphering cell-cell interactions and communication from gene expression. Nat Rev Genet. 2021 Feb;22(2):71–88. doi: 10.1038/s41576-020-00292-x.

3. Bar-Joseph Z, Gitter A, Simon I. Studying and modelling dynamic biological processes using time-series gene expression data. Nat Rev Genet. 2012 Jul 18;13(8):552–64. doi: 10.1038/nrg3244.

4. Baudrimont A, Voegeli S, Viloria EC, Stritt F, Lenon M, Wada T, Jaquet V, Becskei A. Multiplexed gene control reveals rapid mRNA turnover. Sci Adv. 2017 Jul 12;3(7):e1700006. doi: 10.1126/sciadv.1700006.

5. Bernstein JA, Khodursky AB, Lin PH, Lin-Chao S, Cohen SN. Global analysis of mRNA decay and abundance in Escherichia coli at single-gene resolution using two-color fluorescent DNA microarrays. Proc Natl Acad Sci U S A. 2002 Jul 23;99(15):9697–702. doi: 10.1073/pnas.112318199.

6. Blank HM, Perez R, He C, Maitra N, Metz R, Hill J, Lin Y, Johnson CD, Bankaitis VA, Kennedy BK, Aramayo R, Polymenis M. Translational control of lipogenic enzymes in the cell cycle of synchronous, growing yeast cells. EMBO J. 2017 Feb 15;36(4):487–502. doi: 10.15252/embj.201695050

7. Blumberg A, Zhao Y, Huang YF, Dukler N, Rice EJ, Chivu AG, Krumholz K, Danko CG, Siepel A. (2021). Characterizing RNA stability genome-wide through combined analysis of PRO-seq and RNA-seq data. BMC biology, 19(1): 30. https://doi.org/10.1186/s12915-021-00949-x

8. Bolliet V, Labonne J, Olazcuaga L, Panserat S, Seiliez I. Modeling of autophagy-related gene expression dynamics during long term fasting in European eel (*Anguilla anguilla*). Sci Rep. 2017 Dec 20;7(1):17896. doi: 10.1038/s41598-017-18164-6.

9. Cai J, Winkler HH. Identification of tlc and gltA mRNAs and determination of in situ RNA half-life in *Rickettsia prowazekii*. J Bacteriol. 1993 Sep;175(17):5725–7. doi: 10.1128/jb.175.17.5725-5727.1993.

10. Carper D. Deficiency of functional messenger RNA for a developmentally regulated beta-crystallin polypeptide in a hereditary cataract. Science. 1982;217(4558):463-464. doi:10.1126/science.6178163

11. Cho CS, Xi J, Si Y, Park SR, Hsu JE, Kim M, Jun G, Kang HM, Lee JH. Microscopic examination of spatial transcriptome using Seq-Scope. Cell. 2021 Jun 24;184(13):3559–3572.e22. doi: 10.1016/j.cell.2021.05.010.

12. Cho RJ, Campbell MJ, Winzeler EA, Steinmetz L, Conway A, Wodicka L, Wolfsberg TG, Gabrielian AE, Landsman D, Lockhart DJ, Davis RW. A genome-wide transcriptional analysis of the mitotic cell cycle. Mol Cell. 1998 Jul;2(1):65–73. doi: 10.1016/s1097-2765(00)80114-8. PMID: 9702192.

13. Chubb JR, Trcek T, Shenoy SM, Singer RH. Transcriptional pulsing of a developmental gene. Curr Biol. 2006; 16: 1018–1025.

14. Cole SP, Bhardwaj G, Gerlach JH, Mackie JE, Grant CE, Almquist KC, Stewart AJ, Kurz EU, Duncan AM, Deeley RG. Overexpression of a transporter gene in a multidrug-resistant human lung cancer cell line. Science. 1992 Dec 4;258(5088):1650–4. doi: 10.1126/science.1360704.

15. Deana A, Celesnik H, Belasco JG. The bacterial enzyme RppH triggers messenger RNA degradation by 5’ pyrophosphate removal. Nature. 2008 Jan 17;451(7176):355–8.

16. Deng X, Berletch JB, Ma W, Nguyen DK, Hiatt JB, Noble WS, Shendure J, Disteche CM. Mammalian X upregulation is associated with enhanced transcription initiation, RNA half-life, and MOF-mediated H4K16 acetylation. Dev Cell. 2013 Apr 15;25(1):55–68. doi: 10.1016/j.devcel.2013.01.028.

17. DeRisi JL, Iyer VR, Brown PO. Exploring the metabolic and genetic control of gene expression on a genomic scale. Science. 1997 Oct 24;278(5338):680–6. doi: 10.1126/science.278.5338.680.

18. Dölken L, Ruzsics Z, Rädle B, Friedel CC, Zimmer R, Mages J, Hoffmann R, Dickinson P, Forster T, Ghazal P, Koszinowski UH. High-resolution gene expression profiling for simultaneous kinetic parameter analysis of RNA synthesis and decay. RNA. 2008 Sep;14(9):1959–72. doi: 10.1261/rna.1136108.

19. Fisk DG, Ball CA, Dolinski K, Engel SR, Hong EL, Issel-Tarver L, Schwartz K, Sethuraman A, Botstein D, Cherry JM; Saccharomyces Genome Database Project. Saccharomyces cerevisiae S288C genome annotation: a working hypothesis. Yeast. 2006 Sep;23(12):857–65. doi: 10.1002/yea.1400.

20. Furlan M, Galeota E, Gaudio ND, Dassi E, Caselle M, de Pretis S, Pelizzola M. Genome-wide dynamics of RNA synthesis, processing, and degradation without RNA metabolic labeling. Genome Res. 2020 Oct;30(10):1492–1507. doi: 10.1101/gr.260984.120

21. Gaidatzis D, Burger L, Florescu M, Stadler MB. Analysis of intronic and exonic reads in RNA-seq data characterizes transcriptional and post-transcriptional regulation. Nat Biotechnol. 2015 Jul;33(7):722–9. doi: 10.1038/nbt.3269.

22. Galdieri L, Vancura A. Acetyl-CoA carboxylase regulates global histone acetylation. J Biol Chem. 2012 Jul 6;287(28):23865–76. doi: 10.1074/jbc.M112.380519.

23. Gorini L, Maas WK. The potential for the formation of a biosynthetic enzyme in Escherichia coli. Biochim Biophys Acta. 1957 Jul;25(1):208–9. doi: 10.1016/0006-3002(57)90450-x.

24. Gotta SL, Miller OL Jr, French SL. rRNA transcription rate in Escherichia coli. J Bacteriol. 1991 Oct;173(20):6647–9. doi: 10.1128/jb.173.20.6647-6649.1991.

25. Graveley BR, Brooks AN, Carlson JW, Duff MO, Landolin JM, Yang L, Artieri CG, van Baren MJ, Boley N, Booth BW, Brown JB, Cherbas L, Davis CA, Dobin A, Li R, Lin W, Malone JH, Mattiuzzo NR, Miller D, Sturgill D, Tuch BB, Zaleski C, Zhang D, Blanchette M, Dudoit S, Eads B, Green RE, Hammonds A, Jiang L, Kapranov P, Langton L, Perrimon N, Sandler JE, Wan KH, Willingham A, Zhang Y, Zou Y, Andrews J, Bickel PJ, Brenner SE, Brent MR, Cherbas P, Gingeras TR, Hoskins RA, Kaufman TC, Oliver B, Celniker SE. The developmental transcriptome of *Drosophila melanogaster*. Nature. 2011 Mar 24;471(7339):473–9. doi: 10.1038/nature09715.

26. Gray JM, Harmin DA, Boswell SA, Cloonan N, Mullen TE, Ling JJ, Miller N, Kuersten S, Ma YC, McCarroll SA, Grimmond SM, Springer M. SnapShot-Seq: a method for extracting genome-wide, in vivo mRNA dynamics from a single total RNA sample. PLoS One. 2014 Feb 26;9(2):e89673. doi: 10.1371/journal.pone.0089673.

27. Hao N, O’Shea EK. Signal-dependent dynamics of transcription factor translocation controls gene expression. Nat Struct Mol Biol. 2011 Dec 18;19(1):31–9. doi: 10.1038/nsmb.2192.

28. Hawrylycz MJ, Lein ES, Guillozet-Bongaarts AL, Shen EH, Ng L, Miller JA, van de Lagemaat LN, Smith KA, Ebbert A, Riley ZL, Abajian C, Beckmann CF, Bernard A, Bertagnolli D, Boe AF, Cartagena PM, Chakravarty MM, Chapin M, Chong J, Dalley RA, David Daly B, Dang C, Datta S, Dee N, Dolbeare TA, Faber V, Feng D, Fowler DR, Goldy J, Gregor BW, Haradon Z, Haynor DR, Hohmann JG, Horvath S, Howard RE, Jeromin A, Jochim JM, Kinnunen M, Lau C, Lazarz ET, Lee C, Lemon TA, Li L, Li Y, Morris JA, Overly CC, Parker PD, Parry SE, Reding M, Royall JJ, Schulkin J, Sequeira PA, Slaughterbeck CR, Smith SC, Sodt AJ, Sunkin SM, Swanson BE, Vawter MP, Williams D, Wohnoutka P, Zielke HR, Geschwind DH, Hof PR, Smith SM, Koch C, Grant SGN, Jones AR. An anatomically comprehensive atlas of the adult human brain transcriptome. Nature. 2012 Sep 20;489(7416):391–399. doi: 10.1038/nature11405.

29. Houseley J, Tollervey D. The many pathways of RNA degradation. Cell. 2009 Feb 20;136(4):763–76. doi: 10.1016/j.cell.2009.01.019.

30. Huynh-Thu VA, Geurts P. dynGENIE3: dynamical GENIE3 for the inference of gene networks from time series expression data. Sci Rep. 2018 Feb 21;8(1):3384. doi: 10.1038/s41598-018-21715-0.

31. Jao CY, Salic A. Exploring RNA transcription and turnover in vivo by using click chemistry. Proc Natl Acad Sci U S A. 2008 Oct 14;105(41):15779–84. doi: 10.1073/pnas.0808480105.

32. Kalucka J, de Rooij LPMH, Goveia J, Rohlenova K, Dumas SJ, Meta E, Conchinha NV, Taverna F, Teuwen LA, Veys K, García-Caballero M, Khan S, Geldhof V, Sokol L, Chen R, Treps L, Borri M, de Zeeuw P, Dubois C, Karakach TK, Falkenberg KD, Parys M, Yin X, Vinckier S, Du Y, Fenton RA, Schoonjans L, Dewerchin M, Eelen G, Thienpont B, Lin L, Bolund L, Li X, Luo Y, Carmeliet P. Single-Cell Transcriptome Atlas of Murine Endothelial Cells. Cell. 2020 Feb 20;180(4):764–779.e20. doi: 10.1016/j.cell.2020.01.015.

33. Kang HJ, Kawasawa YI, Cheng F, Zhu Y, Xu X, Li M, Sousa AM, Pletikos M, Meyer KA, Sedmak G, Guennel T, Shin Y, Johnson MB, Krsnik Z, Mayer S, Fertuzinhos S, Umlauf S, Lisgo SN, Vortmeyer A, Weinberger DR, Mane S, Hyde TM, Huttner A, Reimers M, Kleinman JE, Sestan N. Spatio-temporal transcriptome of the human brain. Nature. 2011 Oct 26;478(7370):483-9. doi: 10.1038/nature10523

34. Kasturi J, Acharya R, Ramanathan M. An information theoretic approach for analyzing temporal patterns of gene expression. Bioinformatics. 2003 Mar 1;19(4):449–58. doi: 10.1093/bioinformatics/btg020.

35. Kwak H, Fuda NJ, Core LJ, Lis JT. Precise maps of RNA polymerase reveal how promoters direct initiation and pausing. Science. 2013 Feb 22;339(6122):950–3. doi: 10.1126/science.1229386.

36. Kærn M, Elston TC, Blake WJ, Collins JJ. Stochasticity in gene expression: from theories to phenotypes. Nat Rev Genet. 2005 Jun;6(6):451–64. doi: 10.1038/nrg1615.

37. La Manno G, Soldatov R, Zeisel A, Braun E, Hochgerner H, Petukhov V, Lidschreiber K, Kastriti ME, Lönnerberg P, Furlan A, Fan J, Borm LE, Liu Z, van Bruggen D, Guo J, He X, Barker R, Sundström E, Castelo-Branco G, Cramer P, Adameyko I, Linnarsson S, Kharchenko PV. RNA velocity of single cells. Nature. 2018 Aug;560(7719):494–498. doi: 10.1038/s41586-018-0414-6.

38. Lee JS, Nair NU, Dinstag G, Chapman L, Chung Y, Wang K, Sinha S, Cha H, Kim D, Schperberg AV, Srinivasan A, Lazar V, Rubin E, Hwang S, Berger R, Beker T, Ronai Z, Hannenhalli S, Gilbert MR, Kurzrock R, Lee SH, Aldape K, Ruppin E. Synthetic lethality-mediated precision oncology via the tumor transcriptome. Cell. 2021 Apr 29;184(9):2487–2502.e13. doi: 10.1016/j.cell.2021.03.030.

39. Lipshutz RJ, Fodor SP, Gingeras TR, Lockhart DJ. High density synthetic oligonucleotide arrays. Nat Genet. 1999 Jan;21(1 Suppl):20–4. doi: 10.1038/4447.

40. Locke JC, Young JW, Fontes M, Hernández Jiménez MJ, Elowitz MB. Stochastic pulse regulation in bacterial stress response. Science. 2011; 334: 366–369.

41. Mathavan S, Lee SG, Mak A, Miller LD, Murthy KR, Govindarajan KR, Tong Y, Wu YL, Lam SH, Yang H, Ruan Y, Korzh V, Gong Z, Liu ET, Lufkin T. Transcriptome analysis of zebrafish embryogenesis using microarrays. PLoS Genet. 2005 Aug;1(2):260–76. doi: 10.1371/journal.pgen.0010029.

42. Marx V. Tick-tock, it’s RNA o’clock. Nat Methods. 2021 Jun;18(6):597–601. doi: 10.1038/s41592-021-01180-w.

43. Mayer A, di Iulio J, Maleri S, Eser U, Vierstra J, Reynolds A, Sandstrom R, Stamatoyannopoulos JA, Churchman LS. Native elongating transcript sequencing reveals human transcriptional activity at nucleotide resolution. Cell. 2015 Apr 23;161(3):541-554. doi: 10.1016/j.cell.2015.03.010.

44. McManus J, Cheng Z, Vogel C. Next-generation analysis of gene expression regulation–comparing the roles of synthesis and degradation. Molecular BioSystems. 2015; 11: 2680–2689.

45. Meng L, Guo Y, Tang Q, Huang R, Xie Y, Chen X. Metabolic RNA labeling for probing RNA dynamics in bacteria. Nucleic Acids Res. 2020 Dec 16;48(22):12566–12576. doi: 10.1093/nar/gkaa1111.

46. Mortazavi A, Williams BA, McCue K, Schaeffer L, Wold B. Mapping and quantifying mammalian transcriptomes by RNA-Seq. Nat Methods. 2008 Jul;5(7):621–8. doi: 10.1038/nmeth.

47. Mukherjee AS, Beermann W. Synthesis of ribonucleic acid by the X-chromosomes of Drosophila melanogaster and the problem of dosage compensation. Nature. 1965 Aug 14;207(998):785–6. doi: 10.1038/207785a0.

48. Palumbo, M. C., Farina, L., & Paci, P. (2015). Kinetics effects and modeling of mRNA turnover. Wiley interdisciplinary reviews. RNA, 6(3), 327–336. https://doi.org/10.1002/wrna.1277

49. Petersen NS, McLaughlin CS, Nierlich DP. Half life of yeast messenger RNA. Nature. 1976 Mar 4;260(5546):70–2. doi: 10.1038/260070a0.

50. Ranz JM, Castillo-Davis CI, Meiklejohn CD, Hartl DL. Sex-dependent gene expression and evolution of the *Drosophila* transcriptome. Science. 2003 Jun 13;300(5626):1742–5. doi: 10.1126/science.1085881.

51. Rodermund L, Coker H, Oldenkamp R, Wei G, Bowness J, Rajkumar B, Nesterova T, Susano Pinto DM, Schermelleh L, Brockdorff N. Time-resolved structured illumination microscopy reveals key principles of Xist RNA spreading. Science. 2021 Jun 11;372(6547):eabe7500. doi: 10.1126/science.abe7500.

52. Roggenkamp R, Numa S, Schweizer E. Fatty acid-requiring mutant of *Saccharomyces cerevisiae* defective in acetyl-CoA carboxylase. Proc Natl Acad Sci U S A. 1980 Apr;77(4):1814–7. doi: 10.1073/pnas.77.4.1814.

53. Schwalb B, Michel M, Zacher B, Frühauf K, Demel C, Tresch A, Gagneur J, Cramer P. TT-seq maps the human transient transcriptome. Science. 2016 Jun 3;352(6290):1225–8. doi: 10.1126/science.aad9841.

54. Schwalb B, Schulz D, Sun M, Zacher B, Dümcke S, Martin DE, Cramer P, Tresch A. Measurement of genome-wide RNA synthesis and decay rates with Dynamic Transcriptome Analysis (DTA). Bioinformatics. 2012 Mar 15;28(6):884–5. doi: 10.1093/bioinformatics/bts052.

55. Schwanhäusser B, Busse D, Li N, Dittmar G, Schuchhardt J, Wolf J, Chen W, Selbach M. Global quantification of mammalian gene expression control. Nature. 2011 May 19;473(7347):337–42. doi: 10.1038/nature10098.

56. Shah S, Takei Y, Zhou W, Lubeck E, Yun J, Eng CL, Koulena N, Cronin C, Karp C, Liaw EJ, Amin M, Cai L. Dynamics and Spatial Genomics of the Nascent Transcriptome by Intron seqFISH. Cell. 2018 Jul 12;174(2):363–376.e16. doi: 10.1016/j.cell.2018.05.035.

57. Singh J, Padgett RA. Rates of in situ transcription and splicing in large human genes. Nat Struct Mol Biol. 2009 Nov;16(11):1128–33. doi: 10.1038/nsmb.1666.

58. Steiner PA, De Corte D, Geijo J, Mena C, Yokokawa T, Rattei T, Herndl GJ, Sintes E. Highly variable mRNA half-life time within marine bacterial taxa and functional genes. Environ Microbiol. 2019 Oct;21(10):3873–3884. doi: 10.1111/1462-2920.14737.

59. Stavreva DA, Wiench M, John S, Conway-Campbell BL, McKenna MA, Pooley JR, Johnson TA, Voss TC, Lightman SL, Hager GL. Ultradian hormone stimulation induces glucocorticoid receptor-mediated pulses of gene transcription. Nat Cell Biol. 2009 Sep;11(9):1093–102. doi: 10.1038/ncb1922.

60. Stuart GR, Copeland WC, Strand MK. Construction and application of a protein and genetic interaction network (yeast interactome). Nucleic Acids Res. 2009 Apr;37(7):e54. doi: 10.1093/nar/gkp140.

61. Tani H, Mizutani R, Salam KA, Tano K, Ijiri K, Wakamatsu A, Isogai T, Suzuki Y, Akimitsu N. Genome-wide determination of RNA stability reveals hundreds of short-lived noncoding transcripts in mammals. Genome Res. 2012 May;22(5):947–56. doi: 10.1101/gr.130559.111.

62. Wada T, Becskei A. Impact of Methods on the Measurement of mRNA Turnover. Int J Mol Sci. 2017 Dec 15;18(12):2723. doi: 10.3390/ijms18122723.

63. Wagner GP, Kin K, Lynch VJ. Measurement of mRNA abundance using RNA-seq data: RPKM measure is inconsistent among samples. Theory Biosci. 2012 Dec;131(4):281–5. doi: 10.1007/s12064-012-0162-3.

64. Wan Y, Anastasakis DG, Rodriguez J, Palangat M, Gudla P, Zaki G, Tandon M, Pegoraro G, Chow CC, Hafner M, Larson DR. Dynamic imaging of nascent RNA reveals general principles of transcription dynamics and stochastic splice site selection. Cell. 2021 May 27;184(11):2878–2895.e20. doi: 10.1016/j.cell.2021.04.012.

65. Wilusz CJ, Wormington M, Peltz SW. The cap-to-tail guide to mRNA turnover. Nat Rev Mol Cell Biol. 2001 Apr;2(4):237–46. doi: 10.1038/35067025.

66. Wissink EM, Vihervaara A, Tippens ND, Lis JT. Nascent RNA analyses: tracking transcription and its regulation. Nat Rev Genet. 2019 Dec;20(12):705–723. doi: 10.1038/s41576-019-0159-6.

67. Xu Z, Asakawa S. Physiological RNA dynamics in RNA-Seq analysis. Brief Bioinform. 2019 Sep 27;20(5):1725–1733. doi: 10.1093/bib/bby045.

68. Xu Z, Asakawa S. A model explaining mRNA level fluctuations based on activity demands and RNA age. PLoS Comput Biol. 2021 Jul 23;17(7):e1009188. doi: 10.1371/journal.pcbi.1009188.

69. Yamada, T., & Akimitsu, N. (2019). Contributions of regulated transcription and mRNA decay to the dynamics of gene expression. Wiley interdisciplinary reviews. RNA, 10(1), e1508. https://doi.org/10.1002/wrna.1508

70. Yang E, van Nimwegen E, Zavolan M, Rajewsky N, Schroeder M, Magnasco M, Darnell JE Jr. Decay rates of human mRNAs: correlation with functional characteristics and sequence attributes. Genome Res. 2003 Aug;13(8):1863–72. doi: 10.1101/gr.1272403.

71. Yeung KY, Fraley C, Murua A, Raftery AE, Ruzzo WL. Model-based clustering and data transformations for gene expression data. Bioinformatics. 2001 Oct;17(10):977–87. doi: 10.1093/bioinformatics/17.10.977. PMID: 11673243.

72. Zeisel A, Köstler WJ, Molotski N, Tsai JM, Krauthgamer R, Jacob-Hirsch J, Rechavi G, Soen Y, Jung S, Yarden Y, Domany E. Coupled pre-mRNA and mRNA dynamics unveil operational strategies underlying transcriptional responses to stimuli. Mol Syst Biol. 2011 Sep 13;7:529. doi: 10.1038/msb.2011.62.

